# General Anesthesia Activates a Central Anxiolytic Center in the BNST

**DOI:** 10.1101/2023.12.20.572586

**Authors:** Dongye Lu, Seonmi Choi, Jaehong Park, Jiwoo Kim, Shengli Zhao, Camille G. Uldry Lavergne, Quinn Desimone, Bin Chen, Bao-Xia Han, Fan Wang, Nitsan Goldstein

## Abstract

Low doses of general anesthetics like ketamine and dexmedetomidine have anxiolytic properties independent of their sedative effects. How these different drugs exert these anxiolytic effects is not well understood. We discovered a population of GABAergic neurons in the oval division of the bed nucleus of the stria terminalis that is activated by multiple anesthetics and the anxiolytic drug diazepam (ovBNST_GA_). A majority of ovBNST_GA_ neurons express neurotensin receptor 1 (Ntsr1) and innervate brain regions known to regulate anxiety and stress responses. Optogenetic activation ovBNST_GA_ or ovBNST^Ntsr1^ neurons significantly attenuated anxiety-like behaviors in both naïve animals and mice with inflammatory pain, while inhibition of these cells increased anxiety. Notably, activation of these neurons decreased heart rate and increased heart rate variability, suggesting that they reduce anxiety through modulation of the autonomic nervous system. Our study identifies ovBNST_GA_/ovBNST^Ntsr1^ neurons as one of the brain’s endogenous anxiolytic centers and a potential therapeutic target for treating anxiety-related disorders.

**HIGHLIGHTS:**

- General anesthetics and anxiolytics activate a population of neurons in the ovBNST
- Anesthesia-activated ovBNST neurons bidirectionally modulate anxiety-like behavior
- Most anesthesia-activated ovBNST neurons express neurotensin receptor 1
- ovBNST^Ntsr1^ neuron activation shifts autonomic responses to an anxiolytic state

## INTRODUCTION

Anxiety is characterized as a state of high arousal and negative valence, resulting from a person’s or an animal’s enhanced alertness to threatening cues (or internal imagination and reflection of such cues) that are uncertain or temporally/spatially distant^1^. Anxiety disorders, affecting an estimated 14-18% of the population at any time, have become one of the most common psychophysiological disorders^2^. However, common treatments such as selective serotonin reuptake inhibitors (SSRIs) or benzodiazepines are ineffective in many patients or have abuse potential^3,4^. Thus, there is an urgent need for new and effective treatments for anxiety disorders.

Anesthetics and sedatives used in general anesthesia (GA) have demonstrated anxiolytic properties when used at low doses despite fundamental differences in their mechanisms and biochemical properties^5-7^. For example, in humans, sub-anesthesia levels of ketamine, an NMDA receptor antagonist, produces robust, quick-onset, and long-lasting anxiolysis on patients with refractory anxiety disorders or social anxiety disorders^8-10^. Dexmedetomidine, an α2-adrenergic receptor agonist, alleviated anxiety-like behaviors in a rodent model of post-traumatic stress disorder (PTSD)^11^ and in nerve ligation-induced chronic pain^12^, and has long been used to provide peri-operative anxiolysis in animals and human patients^13-15^. Sevoflurane, a GABA-receptor potentiator, also reversed the long-term anxiety-like behaviors caused by a single episode of formalin induced inflammatory pain in rodents^16^. These observations raise the question: how do these drugs that act on different receptors produce the unified outcome of anxiolysis? Is there a common target of GA drugs that mediates their anxiolytic effects?

We have previously hypothesized that some of the effects (e.g. sedation and analgesia) of GA are carried out by GA-activated, rather than suppressed neurons. To this end, we had discovered neurons in the supra optic nucleus that are activated by diverse GA drugs that strongly promote sedation and slow-wave sleep^17^. Similarly, we found a heterogeneous population of GA-activated neurons in the central amygdala (CeA_GA_) that have potent analgesic effects and suppress acute and chronic pain-related behaviors^18^. Interestingly, using the immediate early gene Fos as a marker for recently activated neurons, we also identified a cluster of GA-activated neurons located in the oval division of the bed nucleus of stria terminalis (ovBNST)^18^. The ovBNST is a small nucleus within the anterior BNST, which is often considered part of the extended amygdala. In fact, the anterior BNST and CeA share many common inputs and outputs and have largely identical cell type compositions and molecular expression profiles^19^. Importantly, both human and animal studies suggest that the CeA modulates fear responses while the BNST complex regulates anxiety responses to delayed and/or unpredictable threats^20,21^.

Given the consensus that the anterior BNST is a critical center for anxiety, we hypothesized that GA-activated ovBNST neurons (hereafter referred to as ovBNST_GA_ neurons) have anxiolytic functions. We characterized the anatomical, molecular, and functional profile of ovBNST_GA_ neurons and found that they are sufficient to drive behavioral and autonomic responses characteristic of an anxiolytic state. Together, our results reveal a specific subpopulation of ovBNST neurons as a common mediator of GA-induced anxiolysis that could be a novel target for anxiety therapeutics.

## RESULTS

### Different GA drugs and diazepam all activate a cluster of GABAergic ovBNST neurons

To identify neurons that may underly general anesthetic-induced anxiolysis, we searched for populations that were activated by both anesthetics and anxiolytic drugs (Fig. 1A and 1B). Fos immunostaining revealed a population in the ovBNST (Fig. 1C) that was activated by the anesthetics isoflurane, ketamine, and dexmedetomidine as well as the anxiolytic drug diazepam (a GABA_A_ receptor agonist) (Fig. 1D). Saline and oxygen did not induce considerable Fos expression in this region (Fig. 1D). Though sexual dimorphism has been observed in BNST circuits, we observed comparable numbers of isoflurane-activated ovBNST neurons in brains from male and female mice (Fig. S1). Interestingly, while general anesthetics strongly activate neurons in both the ovBNST and the central amygdala (CeA), diazepam’s effects were weaker in the CeA, suggesting that the ovBNST population (hereafter referred to as ovBNST_GA_) may be unique in its ability to modulate anxiety (Fig. S2).

**Figure 1.**
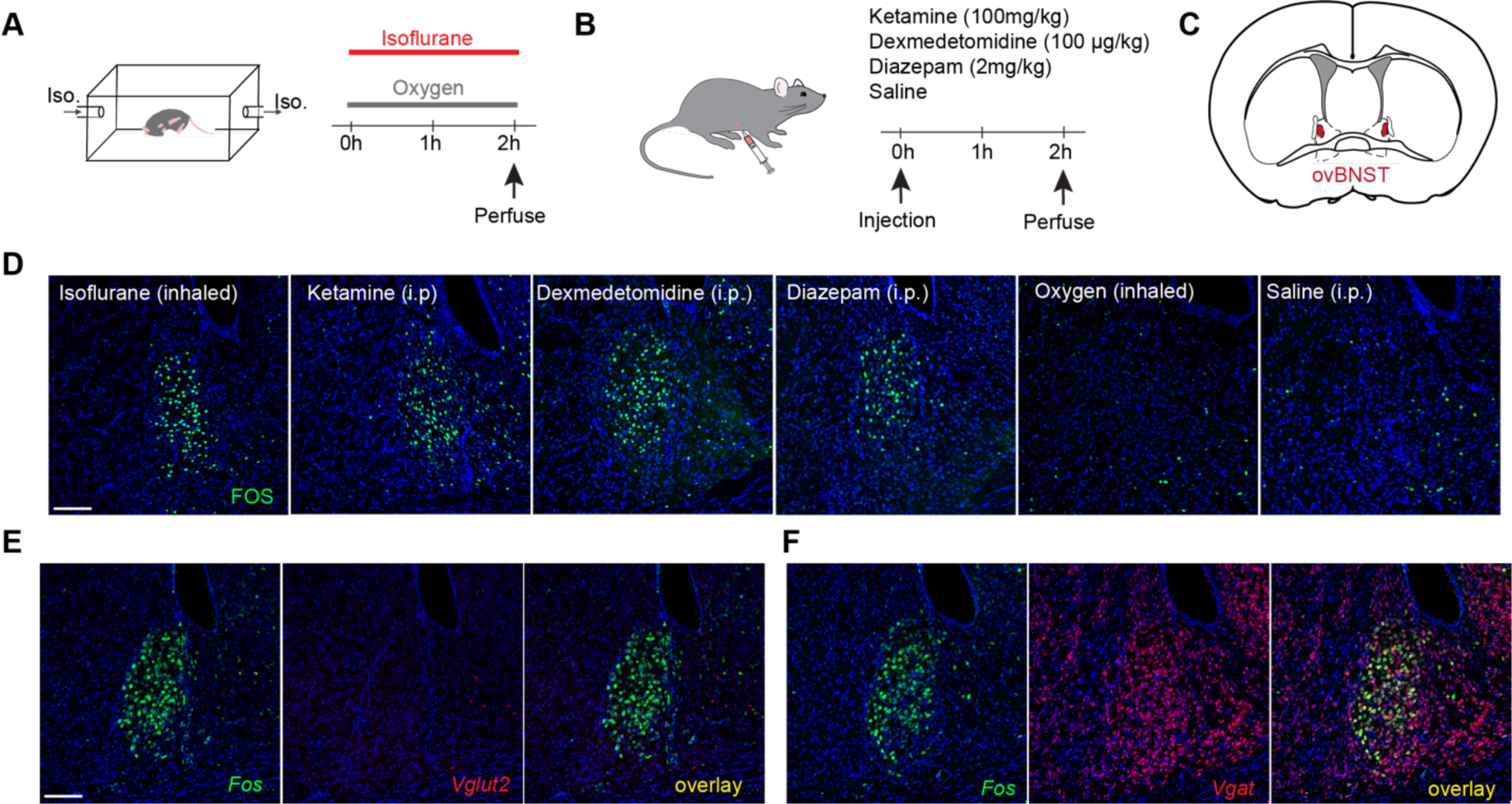
General anesthetics and anxiolytics activate a population of GABAergic neurons in the ovBNST. (**A-B**) Mice were exposed to inhaled 1.5% isoflurane or control oxygen in a chamber (A) or ketamine, dexmedetomidine, diazepam, or control saline intraperitoneally (B) and sacrificed 2 h later to examine Fos expression. (**C**) Representative image showing Fos expression after isoflurane expression in the oval nucleus of the BNST (ovBNST). (**D**) Representative images showing Fos expression in the ovBNST after each stimulus. (**E**) Representative images showing in situ hybridization labeling of *Fos* (green) and *Vglut2* (red) to label excitatory neurons. (**F**) Representative images showing in situ hybridization labeling of *Fos* (green) and *Vgat* (red) to label inhibitory neurons. Scale bar, 100 μm.

The BNST consists many nuclei with both GABAergic and glutamatergic neurons^22^. To determine whether ovBNST_GA_ neurons are predominantly excitatory or inhibitory, we subjected animals to 1 h isoflurane anesthesia and performed *in situ* hybridization for *Fos*, *vGlut2* or *vGat* mRNA as markers for glutamatergic or GABAergic cells. We found very sparse *vGlut2* labeling in the ovBNST and virtually all ovBNST_GA_ neurons were positive for *vGat* and not *vGlut2*. We therefore concluded that ovBNST_GA_ neurons are inhibitory (Fig. 1E and 1F).

### Activity-dependent targeting of ovBNST_GA_ neurons

Isoflurane, ketamine, dexmedetomidine, and diazepam have significantly different mechanisms of action, yet all activated GABAergic neurons in the ovBNST. Are the same neurons activated by these different drugs? We used an activity-dependent method^23^ (Fos-TRAP2) to target ovBNST neurons activated by isoflurane (iso-TRAPed neurons), and subsequently re-exposed mice to different general anesthetics and anxiolytic drugs (Fig. 2A). First, we confirmed efficiency and specificity of the labeling by testing whether iso-TRAPed neurons were reactivated by isoflurane (Fig. 2B). ∼73% of iso-TRAPed neurons (tdTomato+) were also activated (Fos+) by the second exposure to isoflurane and ∼57% of Fos+ neurons were tdTomato+ (Fig. 2D-F). Thus, Fos-TRAP2 is a reasonably efficient and selective tool to label and manipulate ovBNST_GA_ neurons.

**Figure 2.**
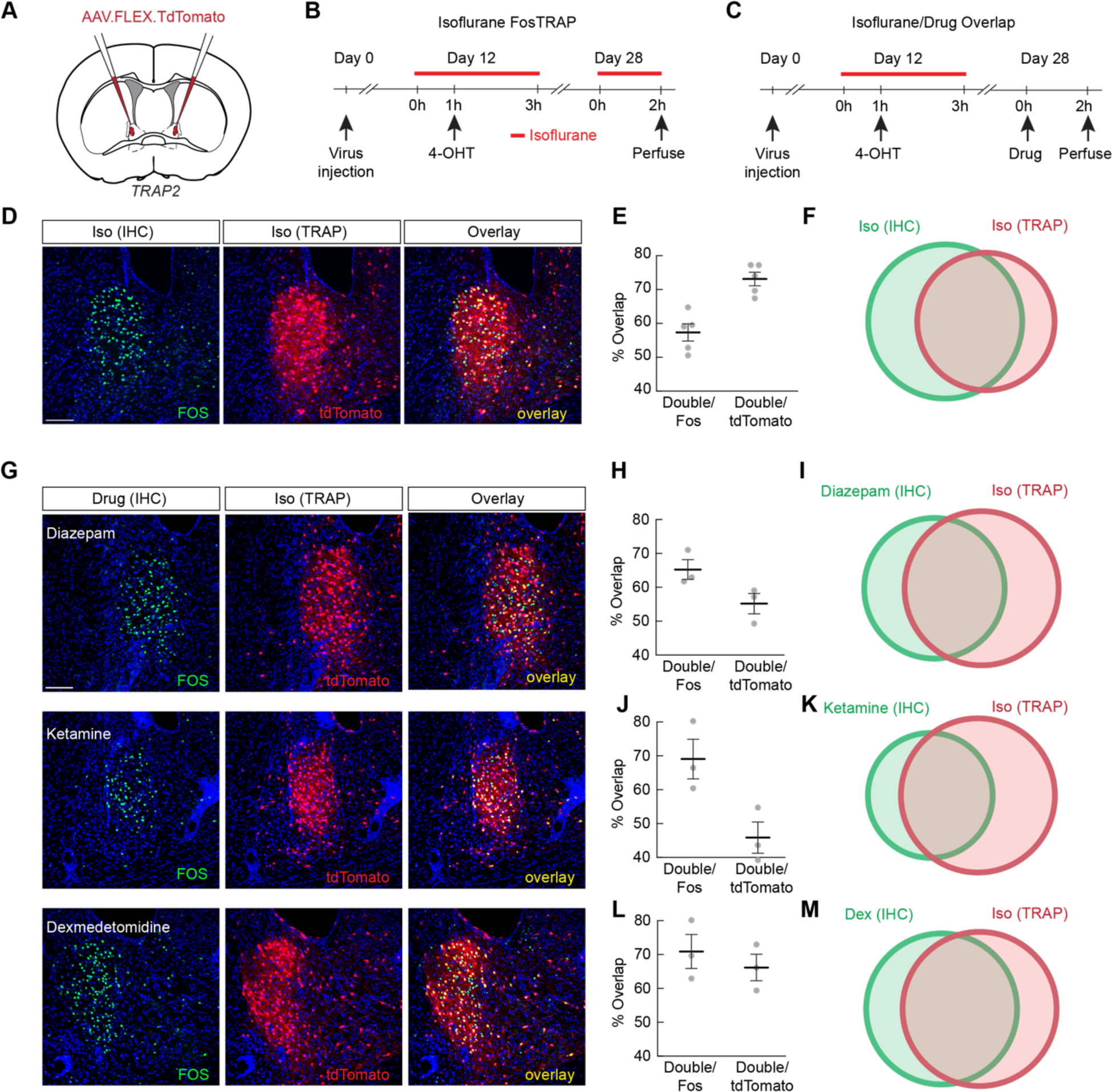
General anesthetics and anxiolytics activate the same subpopulations of ovBNST neurons. (**A**) Fos-TRAP2 mice were injected with Cre-dependent tdTomato in the ovBNST for activity dependent labeling. (**B**) Experimental timeline for validating activity dependent labeling of isoflurane-activated neurons (isoTRAP). (**C**) Experimental timeline for examining overlap between cells activated by isoflurane and by other anesthetic and anxiolytic drugs. (**D**) Representative images showing Fos immunohistochemistry (IHC, green) and isoTRAPed neurons (red) after re-exposure to isoflurane. (**E**) Percent overlap of cells TRAPed by isoflurane exposure and cells expressing Fos after isoflurane re-exposure. (**F**) Schematic representation of average overlaps shown in (E). (**G**) Representative images showing Fos immunohistochemistry and isoTRAPed neurons (red) after exposure to diazepam, ketamine, or dexmedetomidine. (**H**) Percent overlap of cells TRAPed by isoflurane exposure and cells expressing c-Fos after i.p. diazepam. (**I**) Schematic representation of average overlaps shown in (H). (**J**) Percent overlap of cells TRAPed by isoflurane exposure and cells expressing Fos after i.p. ketamine. (**K**) Schematic representation of average overlaps shown in (J). (**L**) Percent overlap of cells TRAPed by isoflurane exposure and cells expressing Fos after i.p. dexmedetomidine. (**M**) Schematic representation of average overlaps shown in (L). Data are depicted as mean ± SEM. Grey dots represent individual mice. Circle areas in Venn diagrams are proportional to the average number of neurons labeled. Scale bar, 100 μm.

Next, iso-TRAPed animals were administered either ketamine, dexmedetomidine, or diazepam, for Fos immunostaining (Fig. 2C). We found that all three drugs were able to activate clusters of ovBNST neurons that considerably overlapped with iso-TRAPed cells (Fig. 2G). Roughly 46% of the iso-TRAPed neurons were activated by ketamine, while ∼69% of the ketamine-activated neurons were iso-TRAPed (Fig. 2G, 2H, and 2I). ∼66% of iso-TRAPed neurons were activated by dexmedetomidine, while ∼71% of the dexmedetomidine-activated neurons were iso-TRAPed (Fig. 2G, 2J, and 2K), and ∼55% of iso-TRAPed neurons were activated by diazepam, while ∼65% of the diazepam-activated neurons were iso-TRAPed (Fig. 2G, 2L, and 2M). Together, these results highlight the existence of a shared population of ovBNST_GA_ neurons activated by diverse anesthetic/anxiolytic drugs.

To rule out the possibility that the Fos expression we observed was a result of stress induced by anesthesia, we subjected mice to either 90 min of restraint, 30 min of random foot shocks, or 10 μL of 4% formalin injected bilaterally into the whisker pad, and subsequently perfused the animals for Fos immunostaining. None of the stressors led to increased Fos expression in the ovBNST, although the neighboring lateral septum (LS), a region implicated in stress responses^24-26^, showed elevated Fos expression in all three conditions (Fig. S3A). We also confirmed the lack of overlap between ovBNST_GA_ and the small number of stress-activated neurons by iso-TRAPing ovBNST_GA_ neurons and re-exposed animals to restraint, foot shocks, or formalin. We found that labeled neurons minimally overlapped with Fos-expressing stress-responsive neurons (Fig. S3B).

### The axonal projection patterns of ovBNST_GA_ neurons

To begin to understand how ovBNST_GA_ neurons integrate into neural circuits involved in modulating anxiety, we examined the axonal projections of ovBNST_GA_ neurons and compared them to previous reports of projection targets from the ovBNST. We injected a Cre-dependent AAV-DIO-GFP virus into the ovBNST and iso-TRAPed these neurons. We found that ovBNST_GA_ neurons densely innervated the anterior-lateral BNST (alBNST) and the ventral BNST (vBNST) (Fig. 3B), both of which are known to project extensively to the paraventricular hypothalamus^27^. We found that, consistent with previous reports, ovBNST_GA_ neurons also densely project to the nucleus accumbens (NAc) and the lateral hypothalamus (LH)^28^ (Fig. 3A and 3D). ovBNST_GA_ neurons sparsely innervate the subthalamic nucleus (STN), the substantia nigra pars compacta or reticulata (SNc/SNr), the periaqueductal grey (PAG), the lateral parabrachial nucleus (lPBN), the reticular formation in hindbrain (Rt), and the medial division of CeA (CeM) (Fig. 3C-H)^29,30^.

**Figure 3.**
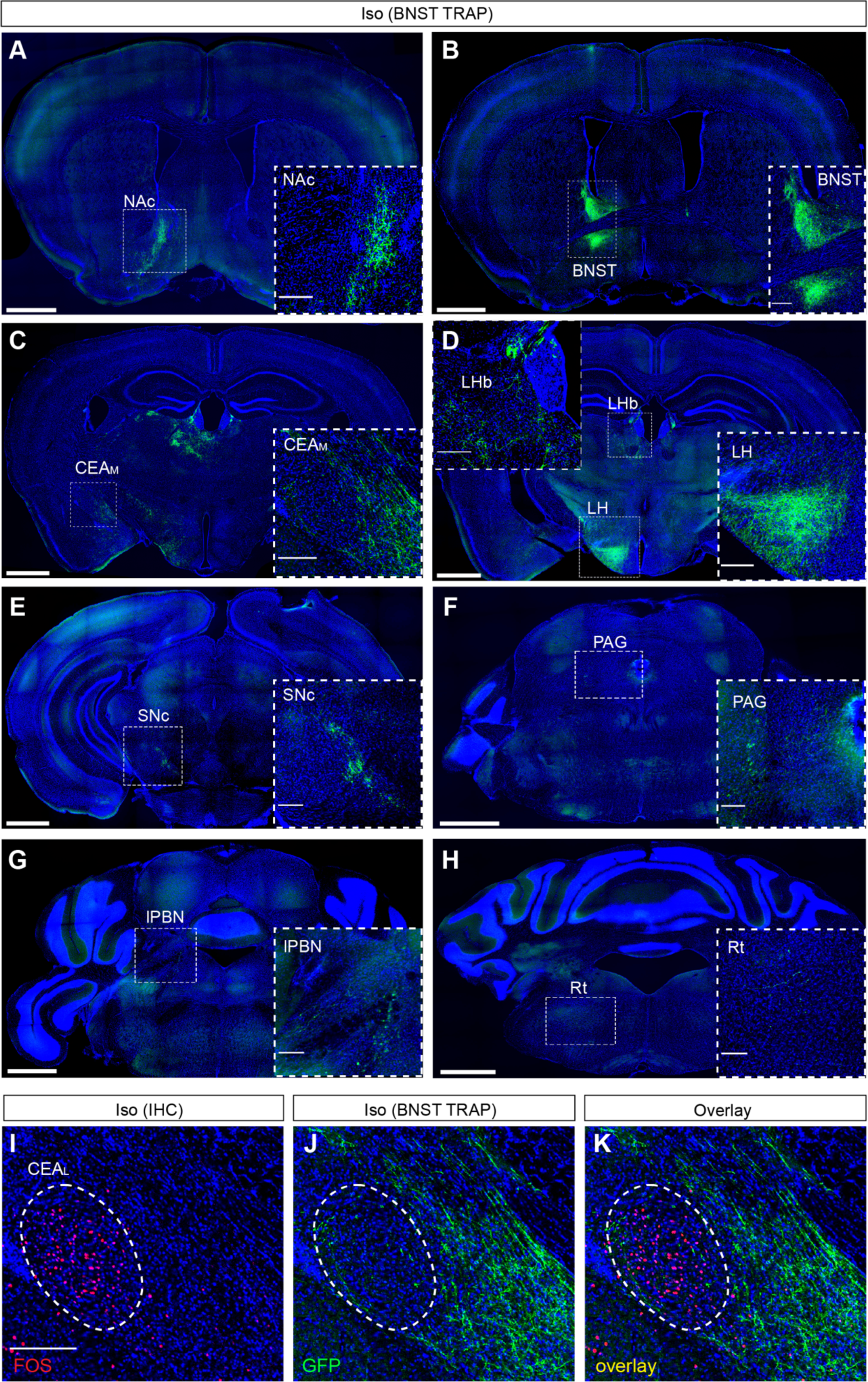
Projection pattern of ovBNST_GA_ neurons. (**A**) Representative image showing ovBNST_GA_ axonal projections to the nucleus accumbens (NAc). Inset, high resolution image of the area inside the white dotted square. (**B**) Representative image showing ovBNST_GA_ cell bodies and axonal projections to the dorsal and ventral regions of the posterior BNST. (**C**) Representative image showing ovBNST_GA_ axonal projections to the central nucleus of the amygdala, medial division (CeA_M_). (**D**) Representative image showing ovBNST_GA_ axonal projections to the lateral habenula (LHb) and lateral hypothalamus (LH). (**E**) Representative image showing sparse ovBNST_GA_ axonal projections to the substantia nigra pars compacta (SNc). (**F**) Representative image showing sparse ovBNST_GA_ axonal projections to ventrolateral periaqueductal grey (vlPAG). (**G**) Representative image showing sparse ovBNST_GA_ axonal projections to the lateral parabrachial nucleus (lPBN). (**H**) Representative image showing sparse ovBNST_GA_ axonal projections to hindbrain reticular nucleus (Rt). (**I-K**) Representative images showing Fos expression in the lateral CeA (CeA_L_, I, red) after isoflurane re-exposure, ovBNST_GA_ axonal projections to the CeA_M_ (J, green), and overlap (K) showing that ovBNST_GA_ neurons do not project to CeA_GA_ neurons. Whole brain images scale bar, 1 mm. High resolution images scale bar, 200 μm.

We noted that ovBNST_GA_ cells do not project to CeA_GA_ cells that we previously found to be strongly analgesic (marked by isoflurane induced Fos expression in the CeA, Fig. 3I-K)^18^. We also observed projections to several areas that have not been previously described including midline thalamic nuclei (paraventricular (PVT) and medial dorsal (MD) nuclei) and the lateral habenula (LHb) (Fig. 3D). Of note, many of the axonal targets of ovBNST_GA_ neurons are known to be involved in modulating stress and anxiety-related behaviors.

### Activation of ovBNST_GA_ neurons reduce anxiety-related but not pain-elicited behaviors

We next investigated the functional role of ovBNST_GA_ neurons. We first asked whether these neurons have pain suppressing effects due to their activation by anesthetics as well as our previous findings that GA-activated CeA_GA_ neurons suppress pain. We bilaterally injected Cre-dependent AAV vectors into the ovBNST of Fos-TRAP2 mice to express either channelrhodopsin (ChR2) for optogenetic activation^31^, enhanced archaerhodopsin (eArch) for optogenetic silencing^32^, or GFP/tdTomato as a control. We then TRAPed ovBNST_GA_ neurons using isoflurane exposure as described above (Fig. 4A). To test how these neurons influence pain related behaviors, we injected formalin unilaterally into the whisker pad and quantified self-caring (face wiping) behaviors with or without laser stimulation (Fig. 4B). Neither activation nor silencing of ovBNST_GA_ neurons had any effect on the amount of time animals spent wiping, suggesting that ovBNST_GA_ neurons, unlike CeA_GA_ neurons, are not analgesic (Fig. 4C-E).

**Figure 4.**
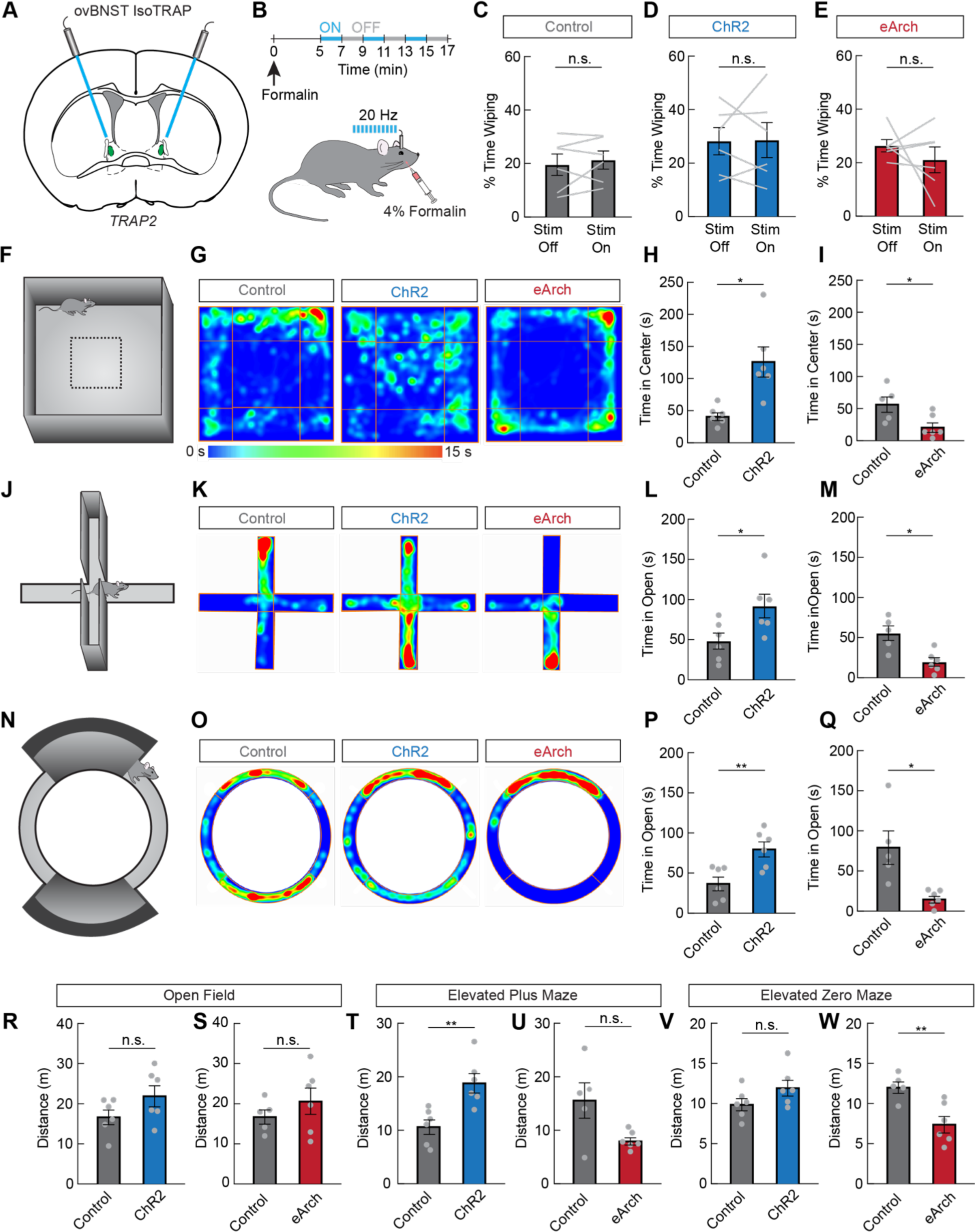
ovBNST_GA_ neurons bidirectionally modulate anxiety-like behavior. (**A**) ovBNST_GA_ neurons were isoTRAPed to express Channelrhodopsin-2 (ChR2), eArchaerhodopsin (eArch), or control GFP bilaterally. Optic fibers were implanted above the injection sites. (**B**) Experimental timeline to test the effects of activating or inhibiting ovBNST_GA_ neurons on acute formalin pain. (**C**) Percent time spent wiping the face during laser off and laser on periods in GFP expressing control mice (n=6, paired samples t-test, n.s.). (**D**) Percent time spent wiping the face during laser off and laser on periods in ChR2 expressing mice (n=6, paired samples t-test, n.s.). (**E**) Percent time spent wiping the face during laser off and laser on periods in eArch expressing mice (n=6, paired samples t-test, n.s.). (**F**) The open field assay was used to assess general locomotion and anxiety-like behavior. (**G**) Representative heat maps showing a mouse expressing control GFP, ChR2, or eArch during the open field assay. Warmer colors indicate more time spent in that area. (**H**) Time spent in the center quarter of the open field in control GFP expressing mice (grey) or ChR2 expressing mice (blue, n=6/group, unpaired t-test, p<0.05). (**I**) Time spent in the center quarter of the open field in control GFP expressing mice (grey) or eArch expressing mice (red, n=5-6/group, unpaired t-test, p<0.05). (**J**) The elevated plus maze was used to assess anxiety-like behavior. (**K**) Representative heat maps showing a mouse expressing control GFP, ChR2, or eArch during the elevated plus maze assay. Warmer colors indicate more time spent in that area. (**L**) Time spent in the open arms of the plus maze in control GFP expressing mice (grey) or ChR2 expressing mice (blue, n=6/group, unpaired t-test, p<0.05). (**M**) Time spent in the open arms of the plus maze in control GFP expressing mice (grey) or eArch expressing mice (red, n=5-6/group, unpaired t-test, p<0.05). (**N**) The elevated zero maze was used to assess anxiety-like behavior. (**O**) Representative heat maps showing a mouse expressing control GFP, ChR2, or eArch during the elevated zero maze assay. Warmer colors indicate more time spent in that area. (**P**) Time spent in the open areas of the zero maze in control GFP expressing mice (grey) or ChR2 expressing mice (blue, n=6/group, unpaired t-test, p<0.01). (**Q**) Time spent in the open areas of the zero maze in control GFP expressing mice (grey) or eArch expressing mice (red, n=5-6/group, unpaired t-test, p<0.05). (**R-W**) Distance traveled during the open field assay (R, n=6/group, unpaired t-test, n.s.; S, n=5-6/group, unpaired t-test, n.s.), elevated plus maze assay (T, n=6/group, unpaired t-test, p<0.01; U, n=5-6/group, unpaired t-test, n.s.), and elevated zero maze (V, n=6/group, unpaired t-test, n.s.; W, n=5-6/group, unpaired t-test, p<0.01). Data are depicted as mean ± SEM. Grey dots and lines represent individual mice. T-tests: n.s., not significant, *p<0.05, **p<0.01.

We next subjected these animals to multiple tests of anxiety-like behavior (open field (OF), elevated plus maze (EPM), and elevated zero maze (EZM)). In rodents, reduced time spent exploring the anxiogenic and open, brightly-lit center zone of the OF and open arms of the EPM and EZM are indicative of increased anxiety-like behaviors^33^. We found that 473 nm laser activation in ChR2-ovBNST_GA_ mice significantly increased the time spent exploring the OF center zone (Fig. 4F-I, 4R, and 4S) and the open arms of the EPM (Fig. 4J-M, 4T and 4U) and EZM (Fig. 4N-Q, 4V and 4W) compared to the control group. By contrast, 561 nm laser stimulation in the eArch-ovBNST_GA_ mice decreased the time spent exploring these anxiogenic areas in all three behavioral tests compared to controls (Fig. 4F-W). Thus, activation of ovBNST_GA_ neurons is anxiolytic, while inhibition is anxiogenic.

Although we did not observe an effect of acute stimulation of ovBNST_GA_ neurons on pain coping behaviors, anxiety and pain are strongly correlated. Having an anxiety disorder is a strong predictor for the development of chronic pain from an acute injury and, conversely, anxiety is a common symptom of those experiencing chronic pain^34,35^. We thus probed whether ChR2 activation of ovBNST_GA_ neurons could also alleviate anxiety-like behaviors in an inflammatory pain model. We used an intraplantar injection of complete Freund’s adjuvant (CFA), which is known to induce long-lasting inflammatory pain and anxiety-like behavior in animals^36,37^. Indeed, activation of ovBNST_GA_ neurons in the CFA model resulted in increased exploration to the OF center and EPM and EZM open arms compared to control CFA-injected mice. Thus, ovBNST_GA_ neurons also reduce anxiety-like behaviors during lasting inflammatory pain (Fig. S4).

### Ntsr1 is a reliable marker for ovBNST_GA_ neurons

Though activity-dependent labeling produced strong behavioral effects when neurons were TRAPed effectively, the success rate of labeling anesthesia-activated ovBNST neurons was very inconsistent. We therefore characterized gene expression in ovBNST_GA_ neurons in order to find a molecular marker. Given the developmental similarities between the CeA and anterior BNST^30^, we first examined marker genes known to be expressed in the CeA including the enzyme *Pkc-delta* (*Pkcd*), the peptides/precursors *Pre-enkephalin* (*Penk1*), *Pre-dynorphin* (*Pdyn*), and *Somatostatin* (*Sst*), and the receptors *Dopamine receptor 1* (*D1r*), *Dopamine receptor 2* (*D2r*), and *Neurotensin receptor 1 (Ntsr1)*. Of the genes tested, the highest overlap with isoflurane activated neurons occurred with cells positive for *Pkcd, Penk1, and Ntsr1*. About ∼69% of isoflurane activated *Fos+* ovBNST neurons expressed *Pkcd*, and ∼32% of *Fos+* neurons expressed *Penk1*. Conversely, ∼51% *Pkcd+* cells and ∼30% *Penk1+* cells were *Fos+* (Fig. S5A and S5B). Notably, ketamine-activated ovBNST neurons had similar proportions of overlap with *Pkcd* and *Penk1* (∼60% and ∼31% of ketamine-induced *Fos+* ovBNST_GA_ neurons expressing *Pkcd* and *Penk1*, respectively, Fig. S5B), consistent with the idea that different GA drugs activated a shared ensemble of ovBNST_GA_ neurons. We also found that a substantial proportion of ovBNST_GA_ neurons express the *Ntsr1* gene encoding the neurotensin receptor type 1 (Fig. S5A). On the other hand, the peptide ligand for Ntsr1, Neurotensin (encoded by *Nts*) is expressed by a small population of ovBNST neurons distinct from ovBNST_GA_ cells (Fig. S5C). Finally, *in situ* hybridization and immunostaining results showed that ovBNST_GA_ neurons did not overlap with neurons expressing *Pdyn* or *Sst* and had very low levels of expression of dopamine receptors *D1r* and *D2r* (Fig. S5C).

While *Pkcd* had the highest degree of overlap with ovBNST_GA_, conflicting results using the Pkcd-Cre BAC-transgenic line in anxiety studies^38-40^ encouraged us to test whether the Ntsr1-Cre driver can be used to target ovBNST_GA_ neurons genetically. We used the knock-in Ntsr1-Cre line^41^ to drive gene expressions in ovBNST_GA_ neurons. We first injected Cre-dependent AAV-DIO-GFP into the ovBNST of Ntsr1-Cre mice (Ntsr1-GFP) and subsequently subjected these mice to isoflurane anesthesia to induce Fos (Fig. 5A and 5B). Overall, ∼71% of Fos+ ovBNST_GA_ neurons were labeled by Ntsr1-GFP, while ∼52% of the Ntsr1-GFP+ ovBNST (ovBNST^Ntsr1^) neurons were Fos+ (Fig. 5C and 5D, Fig. S6A), suggesting that Ntsr1-Cre is a reliable marker for ovBNST_GA_ neurons.

**Figure 5.**
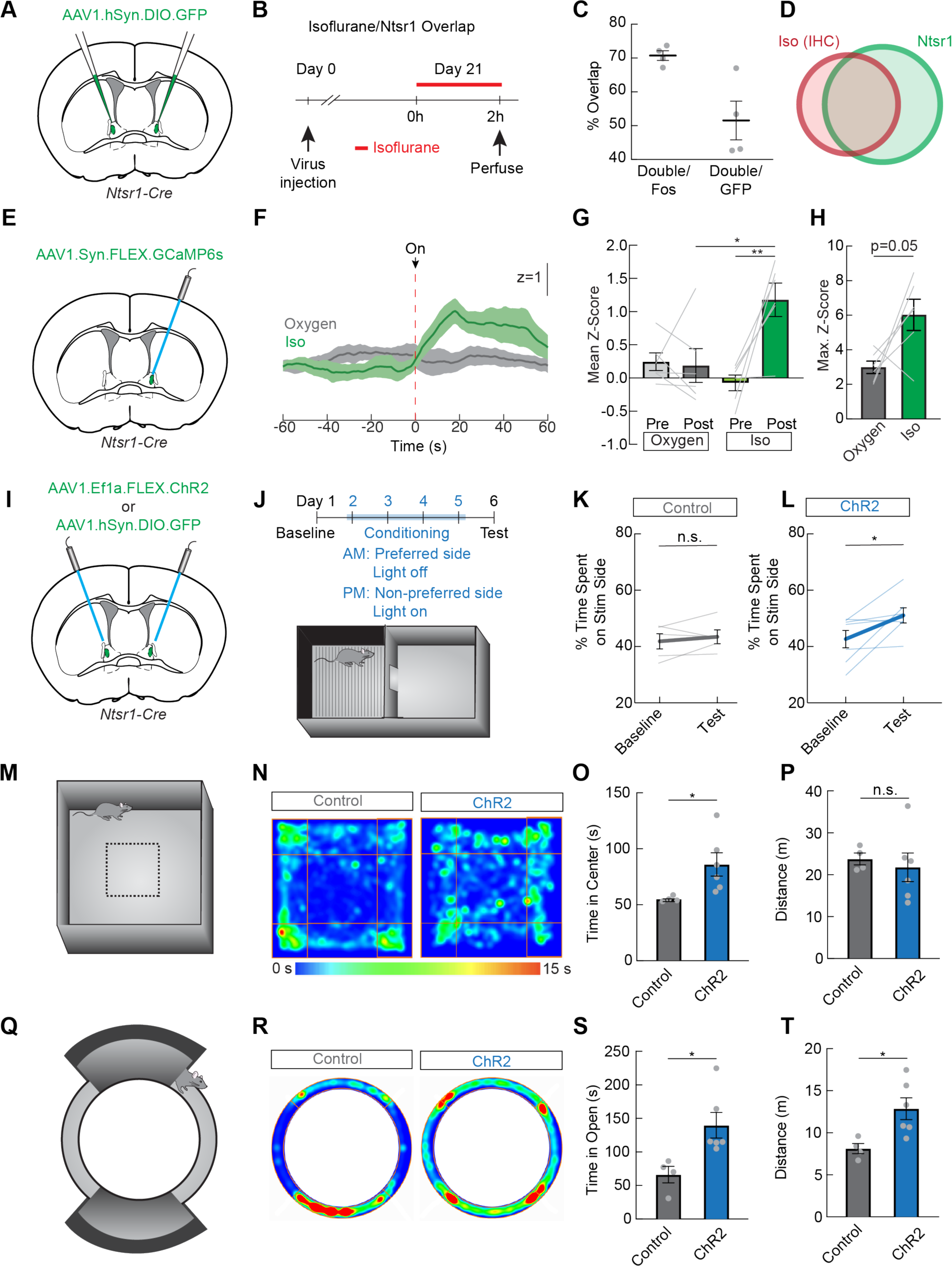
ovBNST^Ntsr1^ neurons are activated by general anesthetics and promote anxiolysis and positive valence. (**A**) GFP was injected into the ovBNST of Ntsr1-Cre mice. (**B**) Experimental timeline to confirm isoflurane activation in Ntsr1-Cre expressing neurons. (**C**) Percent overlap of GFP-positive Ntsr1-Cre cells and cells expressing c-Fos after isoflurane exposure. (**D**) Schematic representation of average overlaps shown in (C). (**E**) GCaMP6s was expressed in the ovBNST of Ntsr1-Cre mice. An optic fiber was implanted above the injection site for dual wavelength fiber photometry recordings in awake, behaving mice. (**F**) Average z-score of GCaMP6s signal in mice exposed to oxygen control (grey) or isoflurane (green). Dark lines represent mean and lighter shaded areas represent SEM. At the time depicted by the dotted red line, during oxygen trials oxygen was turned up from 0.2% to 0.8% (0% isoflurane throughout) and during isoflurane trials isoflurane was turned up from 0% to 2% (0.8% oxygen throughout). (**G**) Mean z-score during fiber photometry experiment shown in (F) (n=6, two-way ANOVA, p<0.05). (**H**) Maximum z-score during oxygen control and isoflurane trials (n=6, paired t-test, p=0.05). (**I**) Control GFP or ChR2 was expressed in Ntsr1-Cre mice for optogenetic activation. (**J**) Conditioned place preference paradigm. Laser stimulation was paired with the less preferred side during conditioning. (**K**) Percent time spent on laser stimulation side in GFP expressing control mice (n=5, paired t-test, n.s.). (**L**) Percent time spent on laser stimulation side in ChR2 expressing mice (n=7, paired t-test, p<0.05). (**M**) The open field assay was used to assess general locomotion and anxiety-like behavior. (**N**) Representative heat maps showing a mouse expressing control GFP or ChR2 during the open field assay. Warmer colors indicate more time spent in that area. (**O**) Time spent in the center quarter of the open field in control GFP expressing mice (grey) or ChR2 expressing mice (blue, n=4-6/group, unpaired t-test, p<0.05). (**P**) Distance traveled during the open field assay in control GFP expressing mice (grey) or ChR2 expressing mice (blue, n=4-6/group, unpaired t-test, n.s.). (**Q**) The elevated zero maze assay was used to assess anxiety-like behavior. (**R**) Representative heat maps showing a mouse expressing control GFP or ChR2 during the elevated zero maze assay. Warmer colors indicate more time spent in that area. (**S**) Time spent in the open areas of the elevated zero maze in control GFP expressing mice (grey) or ChR2 expressing mice (blue, n=4-6/group, unpaired t-test, p<0.05). (**T**) Distance traveled during the elevated zero maze assay in control GFP expressing mice (grey) or ChR2 expressing mice (blue, n=4-6/group, unpaired t-test, p<0.05). Data are depicted as mean ± SEM. Grey dots and lines represent individual mice. T-tests and post-hoc comparisons: n.s., not significant, *p<0.05, **p<0.01.

We also traced axonal projections of ovBNST^Ntsr1^-GFP neurons. The overall pattern of ovBNST^Ntsr1^ neuron projections was similar to that of ovBNST_GA_ neurons. ovBNST^Ntsr1^ neurons’ innervation of many areas, especially in midbrain and hindbrain regions, was denser than that of ovBNST_GA_ neurons, potentially due to the larger number of labeled ovBNST^Ntsr1^ neurons (Fig. S6B-H).

Finally, we used *in vivo* fiber photometry to confirm that ovBNST^Ntsr1^ neurons are activated by general anesthetics. We expressed GCaMP6s in the ovBNST of Ntsr1-Cre mice and implanted an optic fiber to record calcium fluorescence as a proxy for neural activity (Fig. 5E). Isoflurane significantly increased GCaMP6s fluorescence (0.8% O2, isoflurane increased from 0% to 2%) compared to oxygen control trials (0% isoflurane, oxygen increased from 0.2% to 0.8%) (Fig. 5F-H). Taken together, our results demonstrate that Ntsr1 is a reliable marker for most ovBNST_GA_ neurons.

### ovBNST^Ntsr1^ neurons bidirectionally modulate anxiety-like behavior

We next investigated the effect of ovBNST^Ntsr1^ neuron activation on affective and anxiety-like behavior. We expressed ChR2 in ovBNST^Ntsr1^ neurons (Fig. 5I) and subjected mice to a conditioned place preference assay to test whether activation of these neurons promotes positive affect (Fig. 5J). Indeed, mice expressing ChR2, but not control GFP, spent more time in the activation-paired chamber after conditioning, suggesting that ovBNST^Ntsr1^ neuron activation is inherently pleasant (Fig. 5K and 5L). We also found that, like ovBNST_GA_ neuron activation, ChR2 activation of ovBNST^Ntsr1^ neurons led to increased exploration time of the center zone of the OF and the open arms of the EZM (Fig. 5M-T).

Our results show that acutely activating ovBNST^Ntsr1^ neurons reduces anxiety-like behavior and is associated with positive affect. We therefore wondered whether chronically inhibiting these neurons would drive a prolonged anxious state. We expressed the inwardly rectifying potassium channel Kir2.1^42,43^ in ovBNST^Ntsr1^ neurons and measured anxiety-like behavior over a 6 week period (Fig. 6A and 6B). We also validated the inhibition by exposing mice to isoflurane before sacrificing and indeed found reduced c-Fos expression in in the ovBNST in Kir2.1-expressing mice compared to GFP-expressing controls (Fig. 6C and 6D). Consistent with previous reports, time spent in the open arms of the elevated zero maze was relatively stable in control-GFP mice^44^. In Kir2.1-ovBNST^Ntsr1^ mice, however, the time spent in the anxiogenic open arms steadily decreased after viral injection over time, indicative of persistent anxiety (Fig. 6E and 6F). The total distance traveled was not changed in either group, suggesting that locomotion was not significantly affected (Fig. 6G). We also tested mechanical sensitivity using von Frey assays to examine whether a persistently anxious state could cause hypersensitivity. The results showed that withdrawal threshold was unaffected by the expression of Kir2.1 (Fig. 6H-J). These findings show that optogenetic activation of ovBNST^Ntsr1^ neurons acutely attenuates anxiety-like behavior, while long-term inhibition can induce chronic anxiety-like behavior.

**Figure 6.**
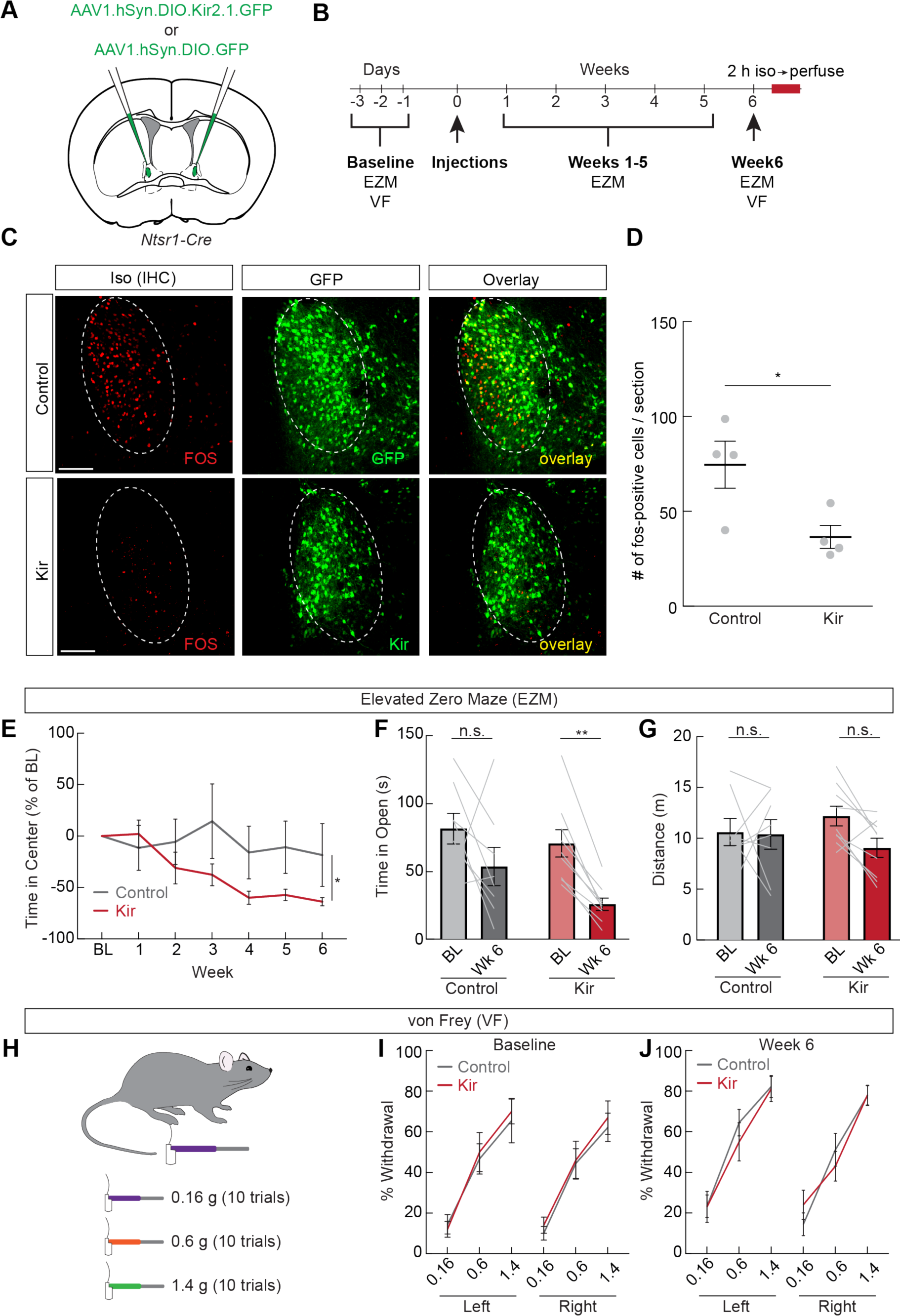
Chronic inactivation of ovBNST^Ntsr1^ neurons drives long term anxiety-like behavior. (**A**) Control AAV-DiO-GFP or AAV-DiO-Kir2.1 was expressed in Ntsr1-Cre mice for chronic inhibition. (**B**) Experimental timeline for chronic inhibition experiments. Elevated zero maze (EZM) and von Frey (VF) tests were performed before and after viral injections in both control and experimental mice. (**C**) Representative images showing isoflurane induced Fos (red) and viral expression (green) in control GFP and Kir2.1 mice. (**D**) Number of Fos positive neurons per section in the ovBNST of control GFP and Kir2.1-expressing mice after isoflurane exposure (n=4/group, unpaired t-test, p<0.05). Grey dots represent an average of 2-3 sections from a single mouse. (**E**) Time spent in the open arms of the EZM (% of baseline, BL) in control (grey) and Kir2.1-expressing (red) mice (n=8-9/group, two-way repeated measures ANOVA, p<0.05). (**F**) Time spent in the open arms of the EZM during the baseline and week 6 tests (n=8-9/group, two-way repeated measures ANOVA, Kir2.1 BL vs. week 6 p<0.01). (**G**) Total distance traveled during the EZM assay during the baseline and week 6 tests (n=8-9/group, two-way repeated measures ANOVA, n.s.). (**H**) The von Frey test was used to determine whether chronic inhibition of ovBNST^Ntsr1^ neurons affects mechanical sensitivity. (**I**) % withdrawal during the von Frey test before viral injections (n=8-9/group, two-way repeated measures ANOVA, n.s.) (**J**) % withdrawal during the von Frey test 6 weeks after viral injections (n=8-9/group, two-way repeated measures ANOVA, n.s.). Data are depicted as mean ± SEM. Grey lines represent individual mice. T-tests, interactions, and post-hoc comparisons: n.s., not significant, *p<0.05, **p<0.01.

### Activation of ovBNST^Ntsr1^ neurons promote autonomic responses characteristic of an anxiolytic state

While these behavioral studies provided evidence for the anxiolytic function of ovBNST_GA_ and ovBNST^Ntsr1^ activations, the predictive validity of rodent behavioral tests for human anxiety is unclear^45^. In humans, anxiety involves characteristic autonomic nervous system (ANS) responses such as increased heart rate and shallow breathing. We therefore decided to measure the effect of ovBNST^Ntsr1^ neurons on heart rate, core body temperature, and breathing.

We implanted telemetric electrocardiogram (ECG) sensors in Ntsr1-Cre mice and expressed either ChR2 or control GFP in ovBNST^Ntsr1^ neurons (Fig. 7A-C). We first tested whether short, continuous activation of ovBNST^Ntsr1^ neurons could affect heart rate by delivering continuous 473 nm light for 5 s while monitoring ECG signals. Consistent with ovBNST^Ntsr1^ neurons’ anxiolytic function, stimulation evoked a significant decrease in heart rate (HR) in ChR2-, but not GFP-expressing mice (Fig. 7D and 7E). We also tested the effects of ovBNST^Ntsr1^ neuron activation over a longer timescale as mice gradually habituated to the stimulation environment (an unfamiliar empty cage). We stimulated ovBNST^Ntsr1^ neurons at 20 Hz using a 3 min on, 3 min off paradigm for 30 mins (Fig. 7F). We found a significant reduction of HR early in the trial when mice were more anxious (Fig. 7G), but the effect of neuronal activation diminished by the final stimulation as animals habituated to the new environment (Fig. 7H and 7I), suggesting the effect of ovBNST^Ntsr1^ is state-dependent.

**Figure 7.**
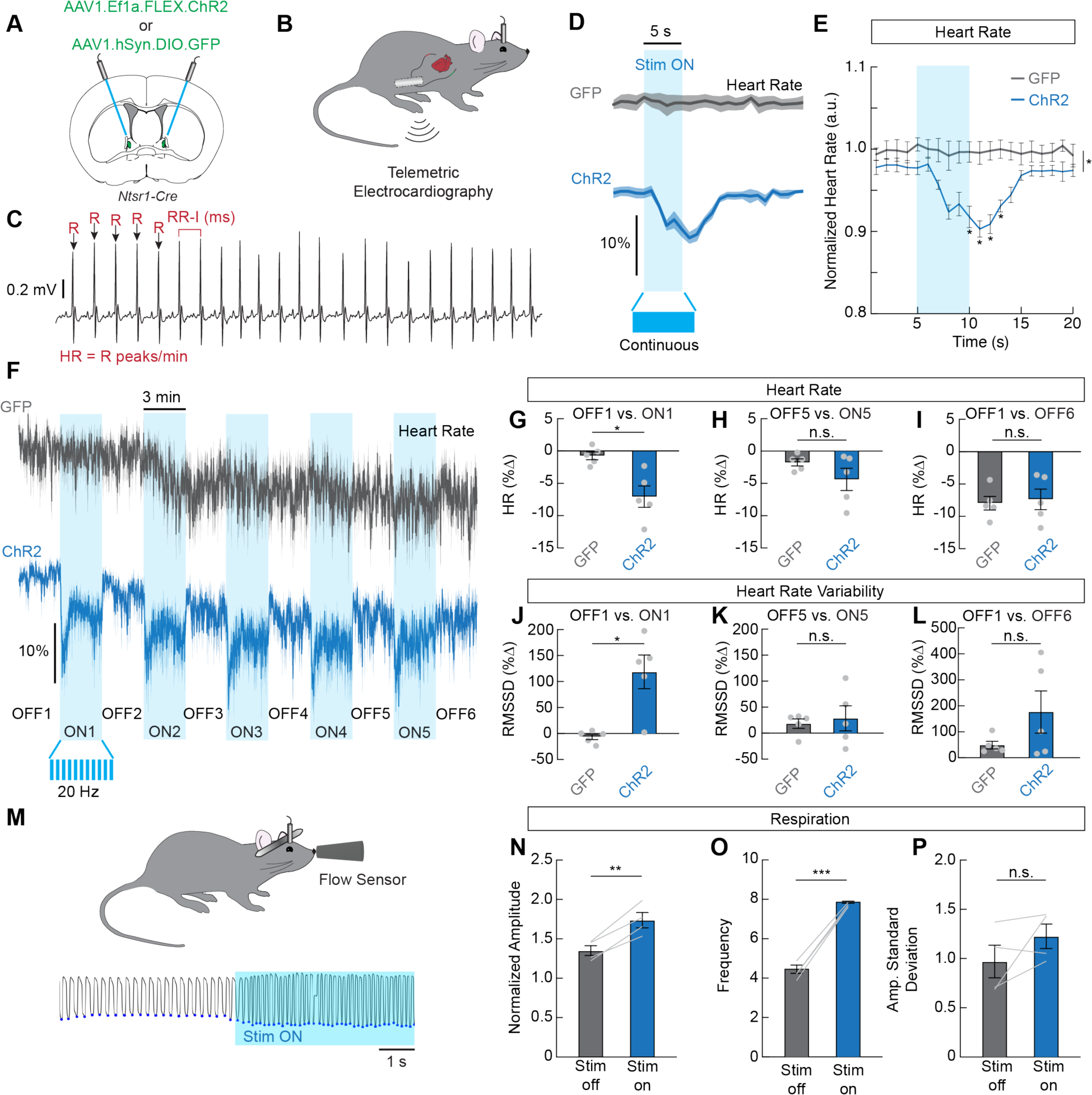
ovBNST^Ntsr1^ neurons shift autonomic output to a less anxious state. (**A**) Control GFP or ChR2 was expressed in Ntsr1-Cre mice for optogenetic activation. (**B**) Wireless telemetry sensors were implanted in the abdomen, with electrodes implanted in the chest. (**C**) Example raw electrocardiography (ECG) trace showing R peaks (arrows). Heart rate was calculated as R peaks/min and heart rate variability was calculated from RR intervals, or the time between successive R peaks. (**D**) Average normalized heart rate in GFP-expressing mice (grey) and ChR2-expressing mice (blue) during a 5 s continuous stimulation with 473 nm laser. Dark lines represent mean and lighter shaded areas represent SEM. (**E**) Mean normalized heart rate from the recordings shown in (D) (n=5/group, repeated measures two-way ANOVA, p<0.001). (**F**) Average normalized heart rate in GFP-expressing mice (grey) and ChR2-expressing mice (blue) during a 30 min trial where 20 Hz 473 nm light was delivered in a 3 min off, 3 min on paradigm. (**G**) Change in heart rate during the first “stim on” period compared to the first “stim off” period in GFP and ChR2 expressing mice (n=5/group, unpaired t-test, p<0.05). (**H**) Change in heart rate during the 5th “stim on” period compared to the fifth “stim off” period in GFP and ChR2 expressing mice (n=5/group, unpaired t-test, n.s.). (**I**) Change in heart rate during the 6^th^ “stim off” period compared to the first “stim off” period in GFP and ChR2 expressing mice (n=5/group, unpaired t-test, n.s.). (**J**) Change in heart rate variability (RMSSD, root mean squared of successive RR intervals) during the first “stim on” period compared to the first “stim off” period in GFP and ChR2 expressing mice (n=5/group, unpaired t-test, p<0.05). (**K**) Change in heart rate variability (RMSSD) during the 5th “stim on” period compared to the fifth “stim off” period in GFP and ChR2 expressing mice (n=5/group, unpaired t-test, n.s.). (**L**) Change in heart rate variability (RMSSD) during the 6^th^ “stim off” period compared to the first “stim off” period in GFP and ChR2 expressing mice (n=5/group, unpaired t-test, n.s.). (**M**) Schematic and representative trace showing breathing during ovBNST^Ntsr1^ neuron stimulation. Blue dots indicate the peaks of inspirations. (**N**) Normalized amplitude during “stim off” periods and “stim on” periods in ChR2-expressing mice (n=4, paired t-test, p<0.01). (**O**) Breathing frequency during “stim off” periods and “stim on” periods in ChR2-expressing mice (n=4, paired t-test, p<0.001). (**P**) Amplitude standard deviation during “stim off” periods and “stim on” periods in ChR2-expressing mice (n=4, paired t-test, n.s.). Data are depicted as mean ± SEM. Grey dots and lines represent individual mice. T-tests, interactions, and post-hoc comparisons: *p<0.05, **p<0.01, ***p<0.001.

A critical autonomic indicator for anxiogenic vs anxiolytic state is heart rate variability (HRV) which reflects the fluctuations of intervals (RR intervals, RRI) between successive heart beats. In healthy individuals, HRV reflects dynamic sympathetic/parasympathetic activity and the body’s capacity to quickly shift the balance in preparation for metabolic and homeostatic needs^46^. In anxiety, however, parasympathetic activity is significant reduced whereas sympathetic activity is elevated, resulting in decreased HRV and a more monotonic heart rate^47^. We analyzed HRV using RMSSD (root mean squared of successive R-R intervals) and compared laser ON vs OFF periods. We found a significant increase in HRV during laser-ON periods in ChR2-ovBNST^Ntsr1^ but not in GFP-ovBNST^Ntsr1^ controls (Fig. 7J). Like the effect on heart rate, the effect of HRV was stronger in the beginning of the trial (Fig. 7J and 7K), while heart rate decreased and HRV increased in both control and experimental animals gradually as mice habituated to the environment (Fig. 7I and 7L). Together, these data suggest that ovBNST^Ntsr1^ neurons could state-dependently regulate autonomic function to suppress anxiety during anxiogenic states.

We also used the 5 s continuous laser stimulation protocol to examine ovBNST^Ntsr1^ neurons’ effects on respiration. We expected that stimulation would decrease breathing rate as slow breathing is associated with calmness, analgesia, anxiolysis, and inhibition of excessive arousal^48-50^. Surprisingly, we observed a significant increase in both amplitude and frequency of breathing that was reliably and precisely triggered by activation of ovBNST^Ntsr1^ neurons (Fig. 7M-P). Notably, the simultaneous increase in amplitude and frequency are distinct from the commonly reported shallow breathing, difficulty in inhalation, and tachypnea in anxiety^51,52^. Since human studies do report increased breathing rates during certain positive emotions (e.g. excitement, happiness)^53^, and exercise increases breathing, the observed increase in respiration amplitude and rate here should not be taken as indicative of negative affect. Finally, we did not observe any consistent, significant effects of ovBNST^Ntsr1^ neurons stimulation on core body temperature (Fig. S7). Overall, our telemetry recordings suggest that ovBNST^Ntsr1^ neurons modulate ANS function by decreasing HR and increasing HRV, thereby shifting the ANS towards a less anxious state.

## DISCUSSION

In this study, we discovered a subpopulation of ovBNST neurons that are commonly activated by diverse general anesthetics and anxiolytics that have different molecular targets. These ovBNST_GA_/ovBNST^Ntsr1^ neurons can state dependently shift both behavioral and ANS responses to a more exploratory, less anxious phenotype. Thus, they represent a key neural substrate through which low-dose general anesthetics and sedatives can induce anxiolysis.

The anterior BNST region has long been known as a major node in the modulation of anxiety. Populations of both anxiogenic and anxiolytic neurons within this region have been described, and it has been suggested that the balance between activity in these populations ultimately determines the overall output of the BNST and its regulation of anxiety^54,55^. Here we show preferential activation of ovBNST_GA_/ovBNST^Ntsr1^ neurons by anxiolytic drugs, but not acute stress, and that this population is strongly anxiolytic. This parallels our previous work that identified a population of anesthesia activated neurons in the CeA, a region that contains both pro- and anti-nociceptive neural populations^56^. CeA neurons located in the capsular division that receive parabrachial input, for example, are hyperactive following injury and are involved in the development of chronic pain^57^. On the contrary, the anesthesia activated CeA_GA_ neurons strongly suppress pain^18^. This raises the possibility that general anesthetics target subpopulations of neurons within GABAergic centers that shift the overall output of these centers to attenuated states of pain or anxiety.

The precise molecular mechanisms through which CeA_GA_ and ovBNST_GA_ neurons are activated by general anesthetics are an interesting topic for future investigation. We also show that diazepam preferentially activates ovBNST_GA_ neurons compared to CeA_GA_ neurons, even though these two sets of cells are known to share embryonic origins and gene expression profiles^30^. Prior studies in acute CeA slices revealed local recurrent inhibitory circuits within the CeA and that diazepam could cause disinhibition of certain populations of CeA neurons^40^ but it is difficult to correlate the concentrations of diazepam used in slice with *in vivo* conditions. Whether the local circuits in the ovBNST further facilitate the disinhibition of ovBNST_GA_ neurons by anesthetics and anxiolytics remains to be carefully examined. Future detailed transcriptomic and connectomic studies of the ovBNST will greatly enable future dissections of this central anxiolytic node and aid the development of potential treatments for anxiety based on activating ovBNST_GA_ neurons. Our finding that ovBNST_GA_ neurons express the neurotensin receptor Ntsr1 supports the possibility that endogenous neurotensin signaling in the ovBNST may regulate anxiety. Small molecule Ntsr1 agonists have been pursued for decades as potential therapeutics for treating conditions such as pain, schizophrenia, obesity, and addiction^58-60^. Our study should further fuel the interest in developing Ntsr1 agonists as treatments for diverse neurologic and psychiatric diseases.

Autonomic responses are an integral component of all emotions including anxiety. The relative ease of collecting autonomic measurements in humans has resulted in a plethora of data on the relationship between physiological responses and various emotional and disease states. For example, generalized anxiety disorder and a number of chronic pain conditions are consistently associated with reduced HRV^47,61,62^. Yet most animal studies rely solely on motor behaviors without measuring autonomic output. Mice are prey animals and are known to hide their behavioral responses to pain and stress and thus measuring these behaviors alone may be insufficient to determine their affective/emotional state. For example, while the opposing anxiolytic and anxiogenic effects of Pkcd+ BNST neurons from different studies^38-40^ may be caused by heterogeneity in the Pkcd population, it is possible that the ambiguity of behavioral readouts in different experimental environments is a factor as well. We argue that adding autonomic response measurement such as HR and HRV will go a long way in improving the validity of pre-clinical animal studies aimed at identifying circuit and molecular targets for treating anxiety and other emotional or affective disorders.

In conclusion, we have identified a population of neurons in the ovBNST that are activated by general anesthetics and anxiolytic drugs. A majority of these neurons express *Ntsr1* and project both locally to other nuclei in the BNST and to brain areas involved in motivation, affect, and autonomic control. Activating these neurons suppresses behavioral and autonomic hallmarks of anxiety and may therefore serve as a therapeutic target in the treatment of anxiety-related disorders.

## ACKNOWLEDGEMENTS

We thank the Wang lab for helpful discussion and feedback throughout the project. This research was funded by the NIH (DE029342 to F.W.), a grant from the Yang-Tan collectives at MIT (to F.W.), and the Jane Coffin Childs Fund for Medical Research (to N.G.).

## AUTHOR CONTRIBUTIONS

D.L. and F.W. initiated the project. D.L., S.C., J.P., J.K., S.Z., C.G.U.L., B.C., Q.D., B.H., and N.G. performed experiments, analyzed data, and/or provided research support. D.L., F.W., and N.G. wrote the manuscript and prepared the figures with comments from all authors.

## STAR* METHODS

### RESOURCE AVAILABILITY

#### Lead Contact

Further information and requests for resources and reagents should be directed to and will be fulfilled by Lead Contact, Fan Wang.

#### Materials Availability

This study did not generate new unique reagents.

#### Data and code availability

All data reported in this paper will be shared by the lead contact upon request. This paper does not report any original code. Any additional information required to reanalyze the data reported in this paper is available from the lead contact upon request.

### EXPERIMENTAL MODEL DETAILS

All experiments were conducted following protocols approved by the Duke University and Massachusetts Institute of Technology (MIT) Institutional Animal Care and Use Committee. Adult (8 to 12 weeks) Fos^2A-iCreER^ (TRAP2) mice (The Jackson Laboratory, strain #030323) and NtsR1^ΔNEO-Cre^ (The Jackson Laboratory, strain #033365) mice were used for experiments. Animals were housed in the vivarium with a 12-hour light/dark schedule with *ad libitum* access to food and water.

### METHOD DETAILS

#### Histology

##### Immunohistochemistry

Mice were transcardially perfused 90-120 min after exposure to a stimulus (anesthetic drug, diazepam, stress, pain) with phosphate buffered saline (PBS, pH 7.4, Thermo Fisher Scientific) followed by 4% paraformaldehyde (PFA). Brains were collected and post-fixed overnight at 4 °C and then transferred to 30% sucrose solution and maintained at 4 °C for 2 days. Brains were frozen in Tissue-Tek O.C.T. (Sakura) then cut into 80 μm thick sections on a cryostat (Leica). Sections were washed with PBS (3 x 5 min) then incubated in 1% Triton-X100 in PBS at room temperature for 2 h, blocked with 10% Blocking One (Nacalai Tesque) in 0.3% Triton-X100 in PBS at room temperature for 1 h, and then incubated at 4 °C in primary antibody solution. The next day sections were washed (3 x 5 min in PBS) and incubated with species specific, minimally cross-reactive secondary antibody solution overnight at 4 °C. Sections were again washed (3 x 5 min in PBS) and counter stained with 4’, 6-diamidino-2-phenylindole (DAPI, Sigma) and mounted with lab-made mounting media^63^.

##### Fluorescence in situ hybridization

Brains were prepared as described above and sectioned at 40 – 60 μm. To determine whether ovBNST_GA_ cells express prodynorphin (Pdyn) or somatostatis (Sst), fluorescence in situ hybridization was performed as described previously^64^. Pdyn, Sst, and Fos probes were used^18^. Fluorescein isothiocyanate-labeled (FITC) Fos probe was used with digoxigenin-labeled (DIG) Pdyn or Sst probes to measure the co-localization with Fos in ovBNST_GA_ cells. To further examine the molecular identities of ovBNST_GA_ cells, HCR in situ hybridization was performed^65^. Briefly, all probes (Glut2, vGat, Drd1, Drd2, Penk1, Pkcd with Fos) were purchased from Molecular Instruments, hybridized with the sections overnight at 37 °C, then amplified with the corresponding fluorescence-tagged (Alexa Fluor 488, 546 or 647) hairpins overnight at 25 °C. Sections were washed, counter stained with DAPI, and mounted on day 3. Co-localization or overlap of each probe was measured against Fos in ovBNST_GA_ cells.

##### Imaging and quantification

Stained sections were imaged with a confocal laser scanning microscope (Zeiss LSM700). Cells were manually counted and quantified using ImageJ (NIH) to co-localize between: (1) *Fos* expressing neurons and neurons expressing the marker of interest and (2) isoflurane activated neurons and neurons activated by a different stimulus. Quantifications were performed by experimenters blinded to experimental conditions.

#### Drugs

Dexmedetomidine (Zoetis, 100 μg/kg), Ketamine (Dechra, 100 mg/kg), and Diazepam (MIT division of comparative medicine, 2 mg/kg) were dissolved in sterile saline and injected intraperitoneally. Formalin (Sigma, 37% formaldehyde) was diluted to 4% in saline and injected unilaterally into the whisker pad (10 μL). Complete Freund’s Adjuvant (CFA, Sigma, 5 mg/mL) was diluted in saline and injected bilaterally into the whisker pad (5 μL each side).

#### IsoTRAP

TRAP2 mice express a tamoxifen-inducible, improved Cre-recombinase estrogen receptor fusion protein (iCreER) from the *Fos* promoter/enhancer elements^23^. To capture general anesthesia-activated neurons in the ovBNST, 2 weeks after stereotaxic deliveries of viruses, TRAP2 animals were anesthetized with 1.5% isoflurane mixed in 0.8 L/min oxygen for 1 hour in a surgery induction chamber to induce the expression of FOS protein and iCreER. Briefly, 4-hydroxy tamoxifen was dissolved in pure ethanol through shaking at room temperature to make a 20 mg/mL solution. The solution was then mixed with corn oil and vacuum centrifuged to form a 10 mg/mL final solution of 4-hydroxy tamoxifen in corn oil. TRAP2 mice were intraperitoneally injected with 4-hydroxy tamoxifen at 50 mg/kg and put back in the induction chamber for 3 more h of isoflurane anesthesia.

#### Surgery

##### Stereotaxic viral injections

We used AAV2/1-EF1a-DIO-hChR2(H134R)-EYFP-WPRE, AAV2/8-CAG-FLEX-EGFP-WPRE, AAV2/8-CAG-FLEX-tdTomato, AAV2/1-hSyn-DIO-EGFP, AAV2-EF1a-DIO-eArch3.0-EYFP, AAV2/1-Syn-Flex-GCaMP6s-WPRE-SV40, and AAV-hSyn-FLEX-loxP-Kir2.1-2A-GFP for this study. To express desired genes in ovBNST neurons in Fos-TRAP2 or *Ntsr1*-Cre animals, mice were briefly anesthetized with isoflurane (3% isoflurane, 0.8L/min oxygen) before being transferred to a stereotaxic frame (David Kopf Instruments) with anesthesia maintained at 1.5% isoflurane mixed in 0.8L/min oxygen. Craniotomies were created with a dental drill (Aseptico) over the ovBNST, whose stereotaxic coordinates relative to bregma were: AP = +0.18 mm, ML = +/- 1.10 mm, DV = -3.65mm (DV coordinates relative to brain surface). The selected virus for the experiment was delivered using glass pipettes at a volume of 200-300 nL and a rate of 60 nL/min.

##### Optic Fiber Implant

For optogenetic activation and inhibition experiments, optic fibers (optogenetics: 200 μm core diameter, 0.22 NA; fiber photometry: 400 μm core, 0.5 NA; RWD) were implanted bilaterally over ovBNST with stereotaxic coordinates: AP = +0.18 mm, ML = +/- 2.23 mm, DV = -3.15 mm (relative to brain surface instead of skull surface) at 20-degree angles. Optic fibers were secured with Metabond (Parkell) and dental cement (Stoelting).

##### Telemetry Sensor Implant

Mice were anesthetized with 1.5% isoflurane mixed in 0.8L/min oxygen. The telemetric transmitters (ETA-F10; Data Sciences International) were implanted intraperitoneally. Briefly, two 1.5 cm midline incisions were made on the lower abdomen through the skin then through the abdominal wall. The transmitter was inserted into the abdominal cavity and secured to the muscle wall. The ECG electrode leads were tunneled through the muscle and under the skin to the right pectoral muscle (negative lead) and the left caudal rib region (positive lead). Muscle and skin openings were closed with sutures and animals were allowed 2 weeks for recovery.

#### Optogenetic Activation and Inhibition

Animals with optic fibers were connected to optical patch cables (0.22NA, 200 μm core diameter; Doric) coupled to a 473 nm (ChR2 activation) or a 561 nm (eArch3.0 inhibition) laser (Opto Engine LLC). Light pulses were generated with Ami-2 Optogenetic interface controlled by ANY-maze. The 473 nm laser was applied in pulses (∼5 mW/mm^2^, 20 Hz, 10 ms pulse width) in anxiety-like behavioral experiments and telemetry experiments and in continuous 5 second bins (∼5 mW/mm^2^) in telemetry experiments. The 561 nm laser was applied continuously (∼10 mW/mm^2^) to animals with eArch3.0 in anxiety-like behaviors.

#### Fiber Photometry

Animals with optic fibers were connected to optical patch cables (0.57NA, 400 μm core diameter; Doric) coupled to a fiber photometry acquisition system (RWD R821). 470 nm and 410 nm light (∼20-40 μW at fiber tip) was delivered through the patch cord. The resulting signal was collected through the same fiber and focused on a detector. Fluorescence was sampled at 90 fps. Mice were placed in a clear plastic cylinder with a small opening for the patch cord and tubing connected to a mobile isoflurane unit. For control oxygen trials, the recording started with a steady flow of 0.2% oxygen and 0% isoflurane. 5 min later, the oxygen was turned up to 0.8%, with isoflurane still at 0%. Recordings proceeded for another 5 min. For isoflurane trials, the recording started with a steady flow of 0.8% oxygen and 0% isoflurane. 5 min later, isoflurane was turned up to 2% and recordings proceeded for another 5 min. Each mouse received one oxygen and one isoflurane trial separated by at least 24 h.

##### Fiber Photometry Analysis

The 410 nm signal was used for motion and baseline correction. Z-scores were calculated using the standard deviation of the entire trace. Mean Z-scores were derived by calculating the average Z-score and dividing by the integration time. Maximum Z-scores were calculated by identifying the highest Z-score value for each mouse between times 0 and 60 s.

#### Behavioral Assays

##### Open field test

To examine anxiety-like behaviors, mice were placed in a 30 * 30 * 30 cm plastic box with white floor and black walls. Each animal was initially placed in the center of the box. Locomotion activities were recorded with ANY-maze (Stoelting) through a webcam (Logitech) for a total of 10 min. Mice received 10 min of laser stimulation throughout the recording session, either 20 Hz pulses with the 473 nm laser for ChR2 and control animals, or continuous light with the 561 nm laser for eArch3.0 and control animals. A 15 * 15 cm center zone was defined in ANY-maze tracking. Distance travelled and time spent in the whole apparatus and in the center zone were recorded and analyzed with ANY-maze.

##### Elevated Plus Maze Test

To examine anxiety-like behaviors, mice were placed in an elevated plus maze for 10 min as described previously^66^. Locomotion activities were recorded with ANY-maze (Stoelting) through a webcam (Logitech). Mice received 10 min of laser stimulation throughout the recording session, either 20 Hz pulses with the 473 nm laser for ChR2 and control animals, or continuous light with the 561 nm laser for eArch3.0 and control animals. The open arms and closed arms were defined in ANY-maze tracking. Distance travelled and time spent in the whole apparatus, in the open arms, and in the closed arms were recorded and analyzed with ANY-maze.

##### Elevated Zero Maze Test

To examine anxiety-like behaviors, mice were placed in an elevated zero maze for 10 min, which, compared to the elevated plus maze, removed the ambiguous and exposed center area between the open arms and closed arms^44^. Locomotion activities were recorded with ANY-maze (Stoelting) through a webcam (Logitech). Mice received 10 min of laser stimulation throughout the recording session, either 20 Hz pulses with the 473 nm laser for ChR2 and control animals, or continuous light with the 561 nm laser for eArch3.0 and control animals. The open arms and closed arms were defined in ANY-maze tracking. Distance travelled and time spent in the whole apparatus, in the open arms, and in the closed arms were recorded and analyzed with ANY-maze.

##### Conditioned Place Preference

Mice were allowed to freely explore a two-sided chamber for 20 min. The two sides were differentiated by the texture of the floor (ribbed vs. smooth) and by the wall pattern (white vs. striped). Animals were tracked using ANY-maze and the time spent in each side was recorded. During the next 4 days, animals were tethered to a patch cord coupled to a 473 nm laser and placed in their preferred side in the morning for 30 min without laser stimulation. In the afternoon, animals were again tethered and placed in their initially non-preferred side for 30 min and optogenetically stimulated as described above. On day 6, animals were again allowed to freely explore the arena for 20 min. The percent time spent in the stimulation-paired side (excluding time spent in the center) before and after conditioning was calculated for each mouse.

##### Stress

To induce stress, mice were placed in a Tailveiner restrainer (Braintree Scientific) for 90 min. For footshock stress, mice were exposed to 180 inescapable electric footshocks (0.4 mA) over a 30 min period. The duration of each shock was pseudo-randomized between 1 to 3 s and the inter-shock intervals were pseudo-randomized between 2 to 25 s.

##### Formalin assay

10 uL of 4% formalin (Sigma) was injected into either the left or right whisker pad of mice to induce acute pain. After the injection, mice displayed 2 distinct phases of self-recuperating behaviors (i.e., grooming/wiping of the injected area): the initial phase that lasted ∼5 min due to acute peripheral pain; and the second phase that lasted more than 30 min due to ongoing inflammation and central sensitization^67^. The grooming/wiping/licking behavior of the injected whisker pad was video recorded immediately after formalin injection. To allow more time for manipulations, optogenetic stimulation was applied to mice at the start of the second phase in 2 min-ON 2 min-OFF cycles for 3 cycles (a total of 12 min, 6 min laser ON, 6 min laser OFF). Laser stimulation was given as 20 Hz pulses with the 473 nm laser for ChR2 and control animals, or continuous light with the 561 nm laser for eArch3.0 and control animals. The fraction of time spent displaying self-recuperating behaviors in each 2 min epoch was rated and calculated by an experimenter blinded to experimental conditions.

##### von Frey Test

Mice were habituated for 2 h to individual plexiglass enclosures atop mesh flooring. On test days, mice were allowed to habituate for 1 h before testing began. Three von Frey filaments were applied (0.16g, 0.6g, and 1.4g) in ascending order. Force was applied until the filament bent. Each filament was applied 10 times to each paw, with at least 30 s in between trials. Withdrawals including lifting, shaking, or licking the paw were recorded, and the percent of trials after which a withdrawal was noted was calculated for each mouse and averaged across groups.

#### Complete Freund’s Adjuvant (CFA)-induced Persistent Inflammation

A single administration of CFA has been reported to induce chronic inflammation pain in mice for over 2 weeks^68^. For each mouse, two 5 uL injections of 1 mg/mL CFA (Sigma) were injected subcutaneously into the bilateral whisker pads. Swelling of injected whisker pads was observed hours after CFA injections. Anxiety-like behavioral tests were performed on day 3, 4, and 5 after CFA injections.

#### Telemetry and Breathing

##### Telemetric Measurements

For 5 s stimulation experiments, mice were placed in an empty cage for 10 min, during which they received 5 s of continuous laser stimulation (∼5 mW/mm^2^ as described above) each minute. For 30 min stimulation experiments, mice were placed in an empty cage for 30 min where they received 20 Hz stimulation in a 3 min off, 3 min on paradigm. Heart rate and the beat-to-beat R-R intervals (RRI) were derived from ECG traces recorded and analyzed with the Ponemah software (Data Sciences International). Heart rate variability (HRV) was calculated using RMSSD as described previously^46^.

##### Breathing

Each mouse was head fixed to a custom elevated running disc setup using a headpost. Mice were tethered to optic fiber patch cables for the delivery of laser and a custom-built airflow sensor (AMW330V, Honeywell) was aligned to the snout of animals. Mice were allowed to habituate on the setup and were tested for 5 min. Mice received 5 s of continuous laser stimulation (∼5mW/mm^2^, with 473 nm laser) during each 30 s trial (5 s of laser followed by 25 s of inter-laser interval). Respiratory activity was measured as previously described^69^. Voltage signals from the sensor were recorded at 250 kHz and down-sampled to 1kHz for analysis.

### QUANTIFICATION AND STATISTICAL ANALYSIS

All statistical analyses were performed in GraphPad Prism 9. Significance levels were indicated as follows: *: P<0.05, **:P<0.01, ***: P<0.001. Sample sizes were determined based on previous publications and common practice for comparable experiments^18,54,70,71^. Descriptive statistical results were presented as mean ± standard error.

Mice were randomly assigned to various treatment groups to receive viruses used for optogenetic manipulations and/or the various stimuli. Data recording and analysis was performed either automatically or by an individual blind to experimental conditions. Incorrectly virally targeted animals and invalid data entries were removed from the study. No sexual dimorphism in either histology or behavioral results was observed in the study, therefore results from males and females were grouped for analysis.

For behavioral experiments, 2-tailed paired or unpaired t tests and repeated measures two-way ANOVAs were used when appropriate with poshoc tests correcting for multiple comparisons (Holm-Šídák). Welch’s correction or the Gessier-Greenhouse correction were used to account for unequal variance. Details on statistical tests and results can be found in Tables S1 and S2.

For telemetric measurements, heart rate, RRI and core body temperature measurements were outputted and logged at 1 Hz. Data entries were removed to exclude the possibility of abnormal consecutive heart beats or incorrect data logging where RRIs changed over 20% for 2 consecutive beats and were higher than 120 ms, as suggested by previous studies^72,73^. This manipulation also ensured basal heart rate data were within the normal range (∼500-700 bpm) for conscious behaving mice^74^. Group data pooling all mice were used for plotting and statistical tests. Two-tailed t tests were performed to compare heart rate and RRI.

