## Supplementary Material for "General Anesthesia Activates a Central Anxiolytic Center in the BNST"

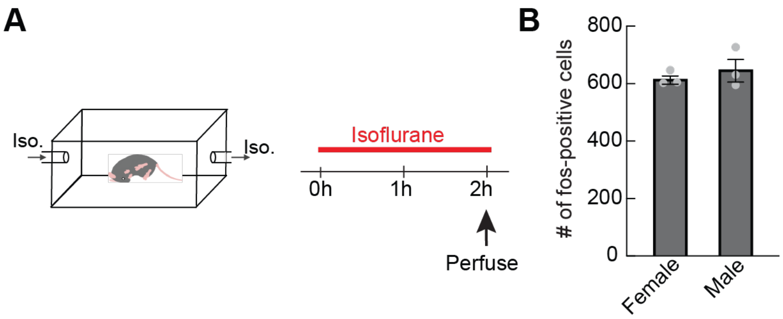


**Figure S1, related to Figure 1. General anesthetics activate ovBNST neurons in male and female mice.** (**A**) Experimental procedure to label isoflurane activated neurons using c-Fos immunohistochemistry. (**B**) Number of Fos positive cells after isoflurane exposure in female and male mice. Data are depicted as mean ± SEM. Grey dots represent individual mice.

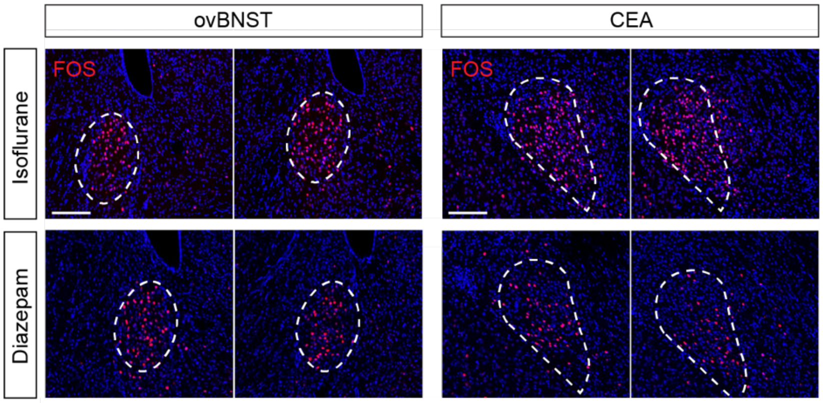


**Figure S2, related to Figure 1. $BNST_{GA}$ neurons are more responsive to anxiolytics than $CEA_{GA}$ neurons.** Representative images showing Fos positive cells in the ovBNST and CEA after isoflurane exposure (top row) and after an injection of diazepam (bottom row). Scale bar, 100 μm.

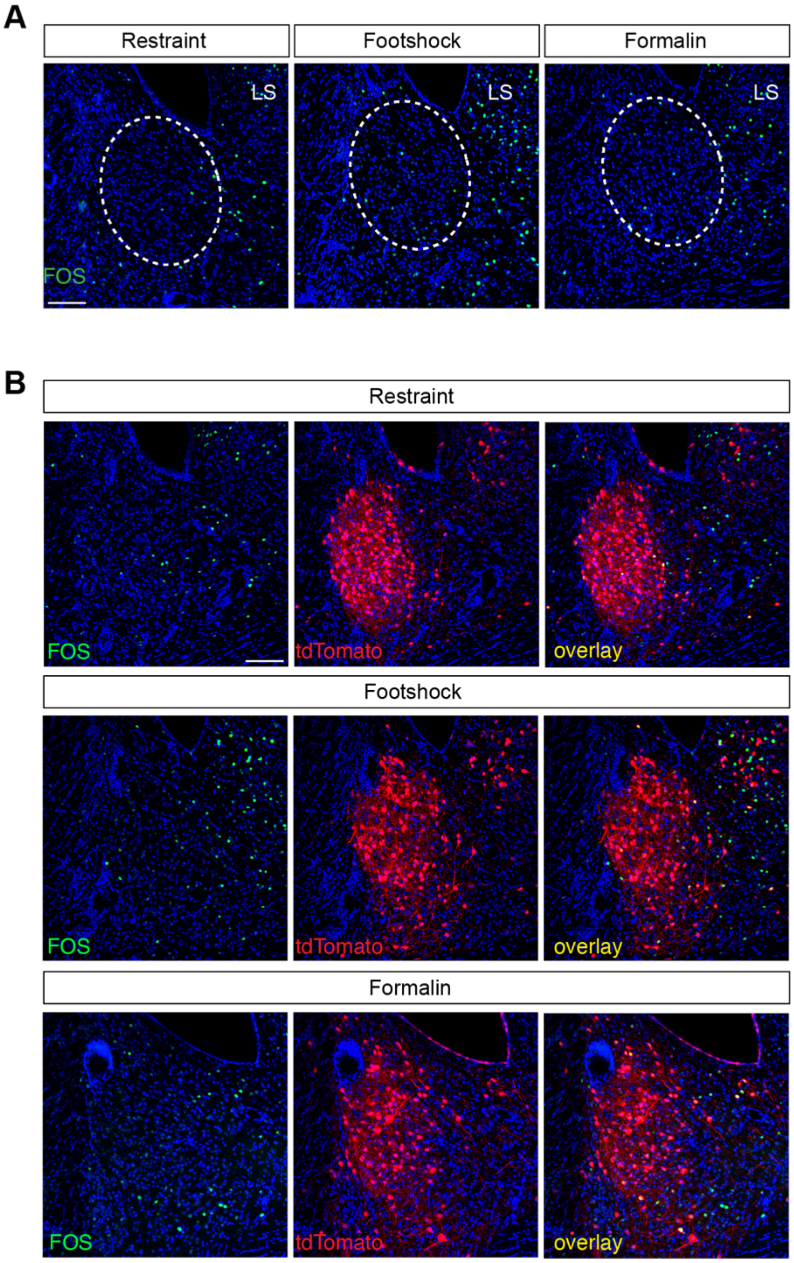


**Figure S3, related to Figure 2. Stress and pain do not activate ovBNST$_{GA}$ neurons.** (**A**) Representative images showing minimal Fos expression in the ovBNST (dotted circle) after stressful or painful stimuli. Fos was expressed in the nearby lateral septum (LS). (**B**) Representative images showing minimal overlap in the ovBNST between isoTRAPed neurons (red) and neurons expressing Fos (green) after exposure to stress or pain. Scale bar, 100 μm.

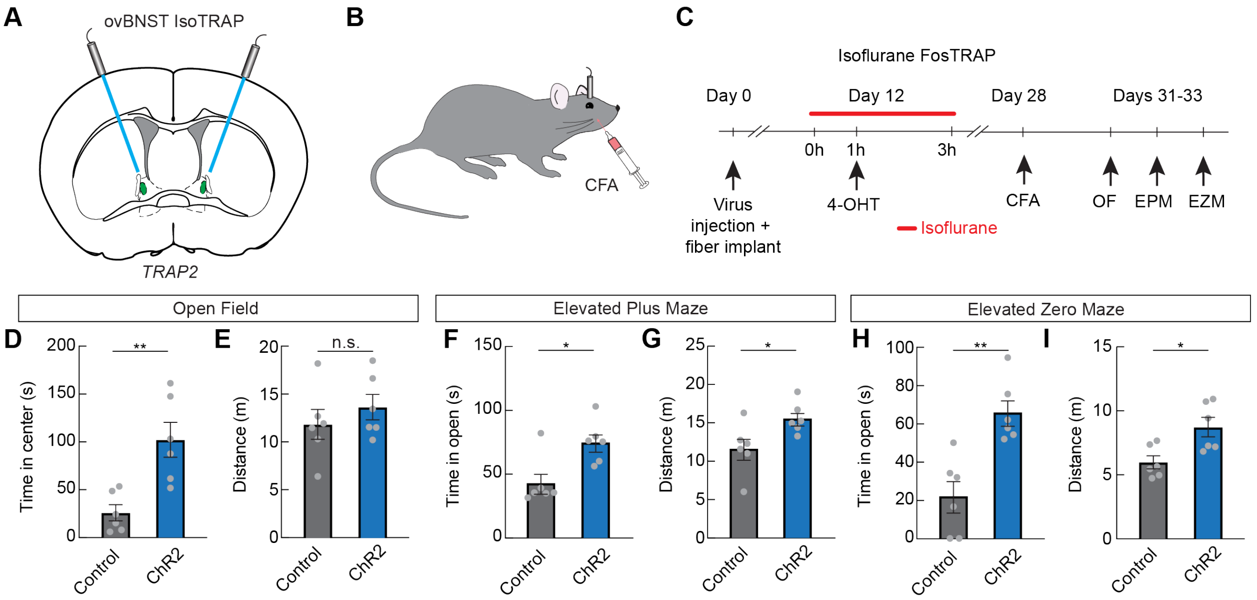


**Figure S4, related to Figure 4. ovBNST$_{GA}$ neurons suppress anxiety-like behavior during lasting inflammatory pain.** (**A**) Control GFP or ChR2 was expressed bilaterally in ovBNST$_{GA}$ neurons. Optic fibers were implanted above injection sites. (**B**) CFA was injected into the whisker pad to evoke persistent inflammation. (**C**) Timeline of experimental procedures to test the effect of ovBNST$_{GA}$ neuron activation on anxiety-like behavior in a model of persistent inflammation. (**D-E**) Time in the center (D) and distance traveled (E) during the open field assay in mice expressing control GFP (grey) or ChR2 (blue, n=6/group, unpaired t-test, $p<0.01$ (D), n.s. (E)). (**F-G**) Time spent in the open arms (F) and distance traveled (G) during the elevated plus maze assay in mice expressing control GFP (grey) or ChR2 (blue, n=6/group, unpaired t-test, $p<0.05$). (**H, I**) Time spent in the open areas (H) and distance traveled (I) during the elevated zero maze assay in mice expressing control GFP (grey) or ChR2 (blue, n=6/group, unpaired t-test, $p<0.01$ (H), $p<0.05$ (I)). Data are depicted as mean ± SEM. Grey dots represent individual mice. T-tests: *$p<0.05$, **$p<0.01$.

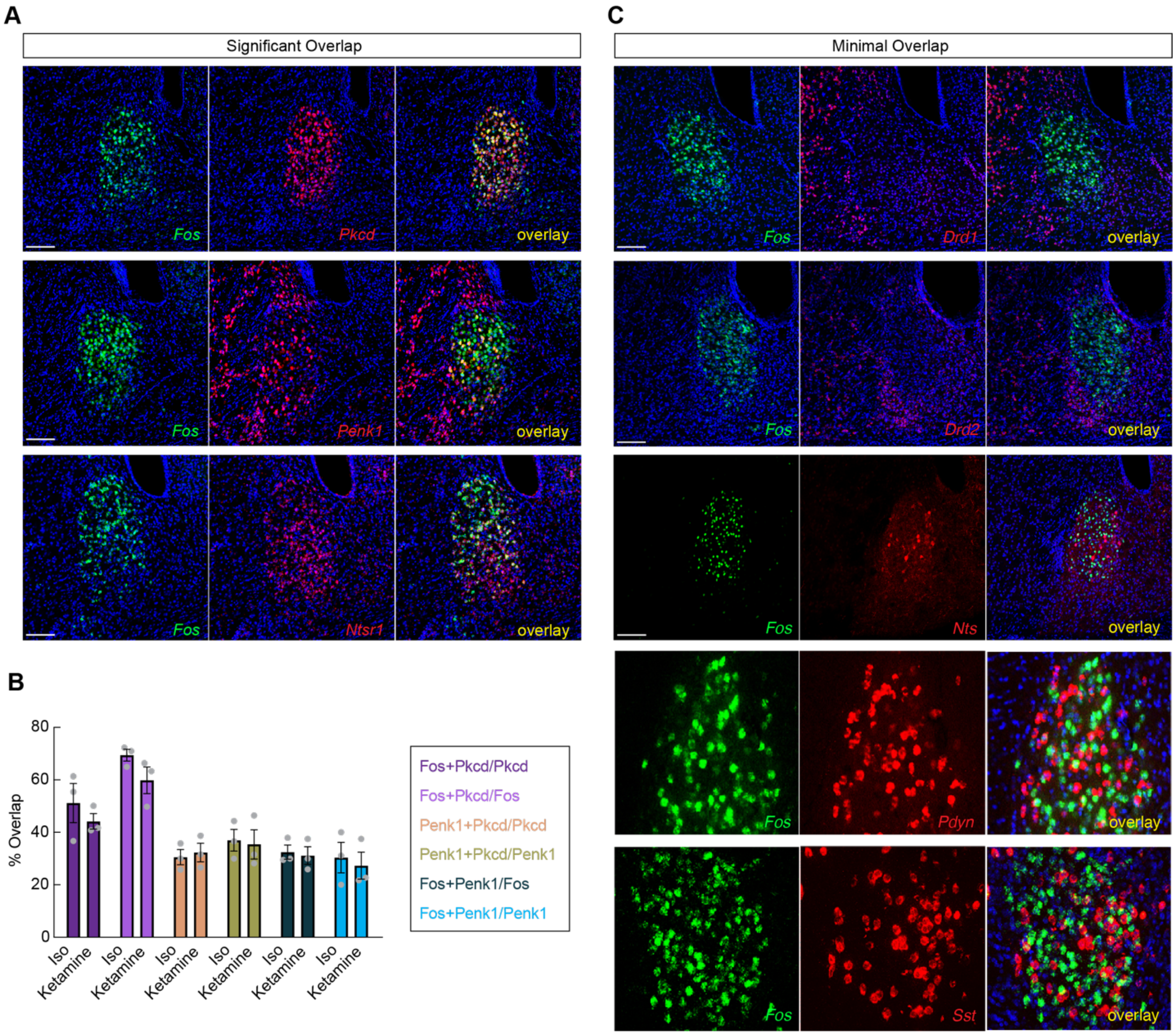


**Figure S5, related to Figure 5. In situ characterization of ovBNST$_{GA}$ neurons.** (**A**) Representative *in situ* hybridization images showing isoflurane-induced *Fos* (green) and three markers (*Pkcd, Penk1,* and *Ntsr1*, red) that overlap with Fos-positive neurons. Scale bar, 100 μm (**B**) % overlap of *Pkcd,* and *Penk1* with isoflurane or ketamine induced *Fos.* (**C**) Representative images showing isoflurane-induced *Fos* (green) and *Drd1, Drd2, Nts, Pdyn,* and *Sst* (red). Scale bar, 100 μm. Data are depicted as mean ± SEM. Grey dots represent individual mice.

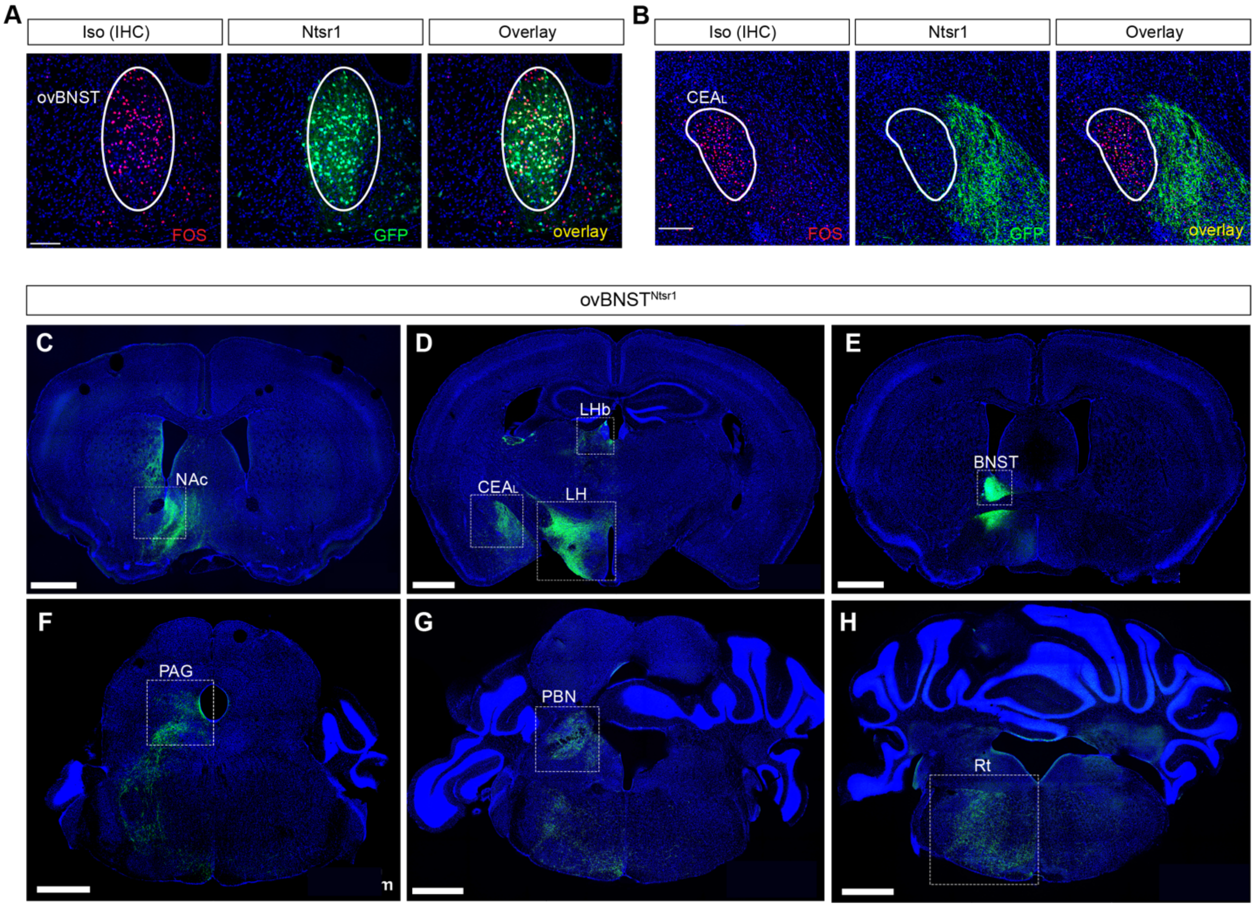


**Figure S6, related to Figure 5. ovBNST$^{Ntsr1}$ neurons are activated by general anesthetics and have similar projection patterns as ovBNST$_{GA}$ neurons.** (**A**) Representative images showing Fos positive neurons after isoflurane exposure (red) and GFP-labeled Ntsr1-Cre neurons (green). Scale bar, 100 μm. (**B**) Representative images showing Fos expression in the lateral CEA (CEA$_L$, I, red) after isoflurane exposure and ovBNST$^{Ntsr1}$ axonal projections to the CEA$_M$ showing that ovBNST$^{Ntsr1}$ neurons do not project to CEA$_{GA}$ neurons. Scale bar, 100 μm. (**C-H**) Representative images showing ovBNST$^{Ntsr1}$ axonal projections to the nucleus accumbens (NAc, C), the CEA$_L$, lateral habenula (LHb), and lateral hypothalamus (LH, D), the BNST (E), the periaqueductal grey (PAG, F), the parabrachial nucleus (PBN, G), and the hindbrain reticular nucleus (Rt, H). Scale bar, 1 mm.

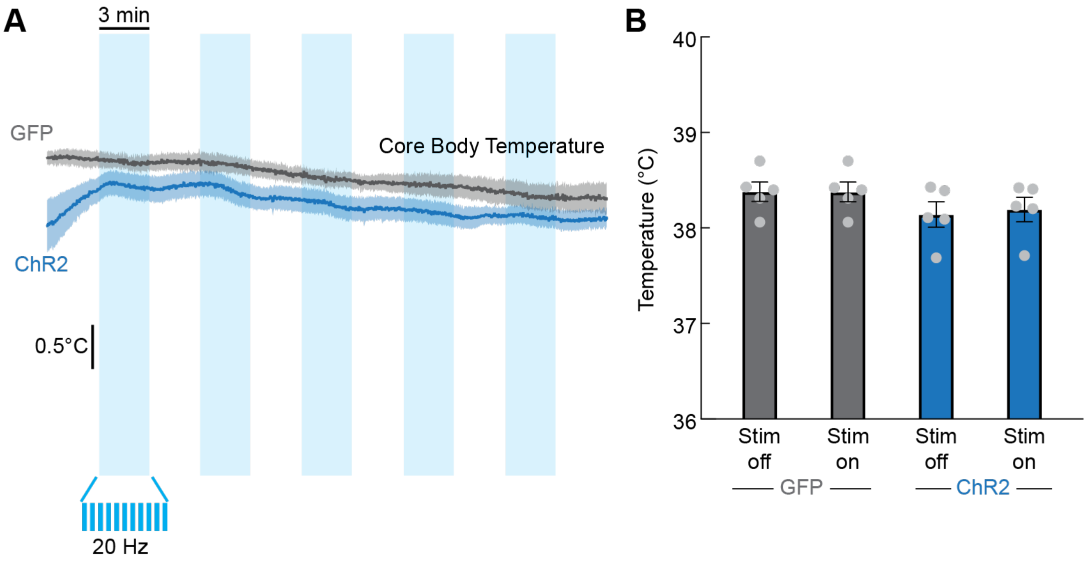


**Figure S7, related to Figure 7. Acute ovBNST$^{Ntsr1}$ neuron activation does not significantly affect core body temperature.** (**A**) Average core body temperature of control GFP-expressing (grey) and ChR2-expressing (blue) mice during 3 min off, 3 min on 20 Hz 473 nm laser stimulation. Dark lines represent mean and lighter shaded areas represent SEM. (**B**) Average core body temperature of control GFP-expressing (grey) and ChR2-expressing (blue) mice during laser off and laser on periods (n=5/group, repeated measures two-way ANOVA, n.s.). Data are depicted as mean ± SEM. Grey dots represent individual mice.

| Figure | Sample Size | Statistical Test | Result |
|---|---|---|---|
| 4C | N = 6 | Paired t-test | p=0.4956 |
| 4D | N = 6 | Paired t-test | p=0.9494 |
| 4E | N = 6 | Paired t-test | p=0.4513 |
| 4H | N = 6/group | Unpaired t-test | *p=0.0137 |
| 4I | N = 5-6/group | Unpaired t-test | *p=0.0398 |
| 4L | N = 6/group | Unpaired t-test | *p=0.0372 |
| 4M | N = 5-6/group | Unpaired t-test | *p=0.0125 |
| 4P | N = 6/group | Unpaired t-test | **p=0.0073 |
| 4Q | N = 5-6/group | Unpaired t-test | *p=0.0349 |
| 4R | N = 6/group | Unpaired t-test | p=0.1375 |
| 4S | N = 5-6/group | Unpaired t-test | p=0.3304 |
| 4T | N = 6/group | Unpaired t-test | **p=0.0054 |
| 4U | N = 5-6/group | Unpaired t-test | p=0.0809 |
| 4V | N = 6/group | Unpaired t-test | p=0.1299 |
| 4W | N = 5-6/group | Unpaired t-test | **p=0.0056 |
| 5G | N = 6 | RM two-way ANOVA<br>Factor 1: iso<br>Factor 2: time<br>Interaction: iso x time<br>Posthoc comparison: Holm-Sidak<br>Oxygen: Pre vs. post<br>Iso: Pre vs. post<br>Pre: oxygen vs. iso<br>Post: oxygen vs. iso | <br>F (1, 5) = 7.182, *p=0.043<br>F (1, 5) = 8.082, *p=0.0361<br>F (1, 5) = 10.82, *p=0.0217<br><br>p=0.8455<br>**p=0.0067<br>p=0.3083<br>*p=0.0170 |
| 5H | N = 6 | Paired t-test | p=0.0548 |
| 5K | N = 5 | Paired t-test | p=0.4825 |
| 5L | N = 7 | Paired t-test | *p=0.0493 |
| 5O | N = 4-6/group | Unpaired t-test | *p=0.0306 |
| 5P | N = 4-6/group | Unpaired t-test | p=0.6081 |
| 5S | N = 4-6/group | Unpaired t-test | *p=0.0123 |
| 5T | N = 4-6/group | Unpaired t-test | *p=0.0132 |
| 6D | N = 4/group | Unpaired t-test | *p=0.0457 |
| 6E | N = 8-9/group | RM two-way ANOVA<br>Factor 1: group<br>Factor 2: time<br>Interaction: group x time | <br>F (1, 15) = 1.906, p=0.1875<br>F (3.276, 49.14) = 3.938, *p=0.0114<br>F (6, 90) = 2.367, *p=0.0361 |
| 6F | N = 8-9/group | RM two-way ANOVA<br>Factor 1: group<br>Factor 2: time<br>Interaction: group x time<br>Posthoc comparison: Holm-Sidak<br>GFP: BL vs. week 6<br>Kir2.1: BL vs. week 6 | <br>F (1, 15) = 3.103, p=0.0985<br>F (1, 15) = 14.43, **p=0.0017<br>F (1, 15) = 0.7749, p=0.3926<br><br>p=0.0633<br>**p=0.0039 |

| | | | |
|---|---|---|---|
| 6G | N = 8-9/group | RM two-way ANOVA<br>Factor 1: group<br>Factor 2: time<br>Interaction: group x time<br>Posthoc comparison: Holm-Sidak<br>GFP: BL vs. week 6<br>Kir2.1: BL vs. week 6 | <br>F (1, 15) = 0.01129, p=0.9168<br>F (1, 15) = 2.175, p=0.1609<br>F (1, 15) = 1.620, p=0.2225<br><br>p=0.8914<br>p=0.0636 |
| 6I | N = 8-9/group | RM two-way ANOVA<br>Factor 1: group<br>Factor 2: filament<br>Interaction: group x filament | <br>F (1, 17) = 0.1357, p=0.7172<br>F (3.830, 65.11) = 35.79, ***p<0.001<br>F (5, 85) = 0.1126, p=0.9893 |
| 6J | N = 8-9/group | RM two-way ANOVA<br>Factor 1: group<br>Factor 2: filament<br>Interaction: group x filament | <br>F (1, 17) = 0.05963, p=0.8100<br>F (4.448, 75.62) = 46.29, ***p<0.001<br>F (5, 85) = 0.7642, p=0.5781 |
| 7E | N = 5/group | RM two-way ANOVA<br>Factor 1: group<br>Factor 2: time<br>Interaction: group x time<br>Posthoc comparison: Holm-Sidak<br>0-1s: GFP vs. ChR2<br>1-2s: GFP vs. ChR2<br>2-3s: GFP vs. ChR2<br>3-4s: GFP vs. ChR2<br>4-5s: GFP vs. ChR2<br>5-6s: GFP vs. ChR2<br>6-7s: GFP vs. ChR2<br>7-8s: GFP vs. ChR2<br>8-9s: GFP vs. ChR2<br>9-10s: GFP vs. ChR2<br>10-11s: GFP vs. ChR2<br>11-12s: GFP vs. ChR2<br>12-13s: GFP vs. ChR2<br>13-14s: GFP vs. ChR2<br>14-15s: GFP vs. ChR2<br>15-16s: GFP vs. ChR2<br>16-17s: GFP vs. ChR2<br>17-18s: GFP vs. ChR2<br>18-19s: GFP vs. ChR2<br>19-20s: GFP vs. ChR2 | <br>F (1, 8) = 10.11, *p=0.0130<br>F (3.303, 26.42) = 11.62, ***p<0.001<br>F (19, 152) = 10.54, ***p<0.001<br><br>p=0.7472<br>p=0.7472<br>p=0.7472<br>p=0.7472<br>p=0.3809<br>p=0.7472<br>p=0.3809<br>p=0.1828<br>p=0.1718<br>*p=0.0403<br>*p=0.0162<br>*p=0.0105<br>*p=0.0239<br>p=0.0898<br>p=0.7425<br>p=0.7425<br>p=0.7425<br>p=0.7425<br>p=0.4939<br>p=0.7472 |
| 7G | N = 5/group | Unpaired t-test | *p=0.0155 |
| 7H | N = 5/group | Unpaired t-test | p=0.2067 |
| 7I | N = 5/group | Unpaired t-test | p=0.7633 |
| 7J | N = 5/group | Unpaired t-test | *p=0.0169 |
| 7K | N = 5/group | Unpaired t-test | p=0.7093 |
| 7L | N = 5/group | Unpaired t-test | p=0.1927 |
| 7N | N = 4 | Paired t-test | **p=0.0067 |
| 7O | N = 4 | Paired t-test | ***p=0.0002 |
| 7P | N = 4 | Paired t-test | p=0.2433 |

**Table S1. Main figures statistical tests and results.**

| Figure | Sample Size | Statistical Test | Result |
|---|---|---|---|
| S4D | N = 6/group | Unpaired t-test | **p=0.0066 |
| S4E | N = 6/group | Unpaired t-test | p=0.4002 |
| S4F | N = 6/group | Unpaired t-test | *p=0.0120 |
| S4G | N = 6/group | Unpaired t-test | *p=0.0368 |
| S4H | N = 6/group | Unpaired t-test | **p=0.0022 |
| S4I | N = 6/group | Unpaired t-test | *p=0.0154 |
| S7B | N = 5/group | RM two-way ANOVA<br>Factor 1: group<br>Factor 2: stimulation<br>Interaction: group x stimulation<br>Posthoc comparison: Holm-Sidak<br>GFP: Stim off vs. stim on<br>ChR2: Stim off vs. stim on | <br>F (1, 8) = 1.614, p=0.2396<br>F (1, 8) = 3.582, p=0.0951<br>F (1, 8) = 4.178, p=0.0752<br><br>p=0.9174<br>*p=0.0238 |

**Table S2. Supplemental figures statistical tests and results.**
